# DIYNAFLUOR: An Affordable DIY Plug-and-Play Nucleic Acid Fluorometer for eDNA Quantification in Resource Limited Settings

**DOI:** 10.1101/2024.12.16.626200

**Authors:** Will Anderson, Fiach Antaw, Sophie Kenny, Harshita Rupani, Ramlah Khamis, Nicolas Constantin, Vinay Kumar, Anna Gemmell, Craig Bell, Matt Trau, Darren Korbie

## Abstract

Nucleic Acid (NA) fluorometry is widely employed for quantifying environmental DNA (eDNA) samples and their downstream DNA sequencing libraries, owing to its sensitivity, accuracy, and speed. However, the high cost of NA fluorometers presents a barrier to eDNA sequencing in resource limited settings (RLSs). For instance, at ∼$1.5-3.3k USD, current NA fluorometers present a greater capital cost than ONT’s $1k USD portable MinION third-generation Nanopore sequencing platform. The collapse of international scientific device and consumable supply chains during the COVID-19 pandemic also highlighted the need for distributed manufacturing of molecular research tools to mitigate the impact of increased pricing to RLSs. To address these challenges, we have developed the “DIYNAFLUOR” (DIY Nucleic Acid FLUORometer), a portable, open-source, <$40 USD NA fluorometer, designed using readily available off-the-shelf components, simple 3D-printed parts, and plug-and-play, solder-free assembly. The DIYNAFLUOR was primarily designed to be compatible with the popular DNA-centric Qubit High Sensitivity (HS) and Broad Range (BR) assay kits. Notably, the DIYNAFLUOR demonstrated an ‘in-assay’ Limit of Detection with the Qubit HS kit of 0.0028 ng/μL, and an average absolute bias of 0.018 ng/μL across a 0–10 ng/μL working range using a 2-point linear calibration methodology. Device verification was performed by comparative measurements with a Qubit 4 fluorometer in a busy biotechnology laboratory, and build instructions were validated through assembly and qualification of three DIYNAFLUOR devices by researchers outside the primary design team. We also describe a custom “extreme”low-cost assay, <13¢ USD per-measurement, that uses SYBR Safe dye to quantify DNA across a working range of 0-0.5 ng/μL. This assay reports a lower sensitivity and accuracy than commercial kits but may be of use to RLSs in times of extreme resource constraints or as a teaching tool for STEM educators. To demonstrate its practical application for field-based eDNA analysis in RLSs, the DIYNAFLUOR was used to perform all quality control measurements throughout the preparation of a 16S and 18S metabarcoding library generated from eDNA extracted from Australian lake water, leading to the successful identification of Australian fauna via Nanopore sequencing. Finally, configurations of the DIYNAFLUOR for RNA and Protein quantification are briefly described.

**TOC:** 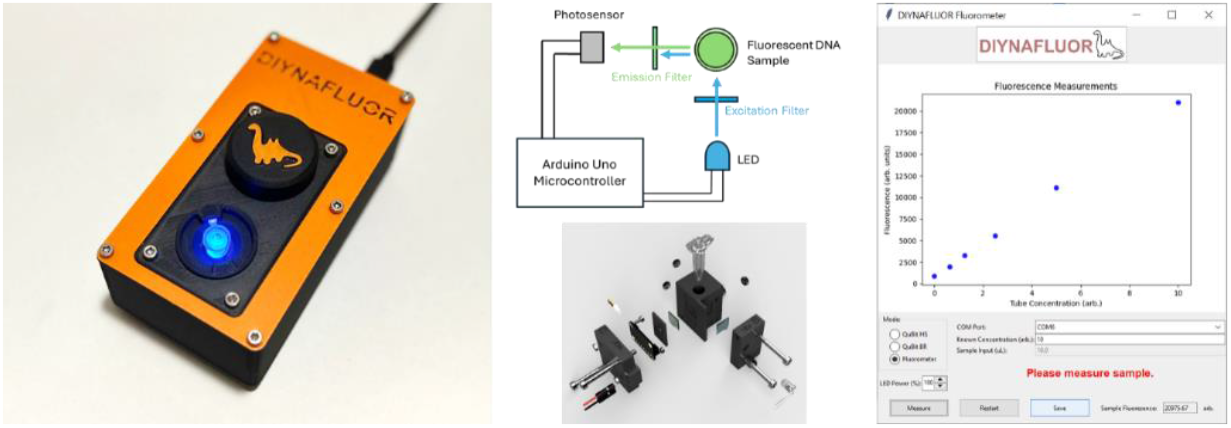

## Introduction

Environmental DNA (eDNA) analysis in Resource Limited Settings (RLSs), i.e., settings where access to research infrastructure is limited to basic resources, has garnered considerable interest in recent years due in part to the increased accessibility afforded by Nanopore third-generation sequencing [1]. This technology has demonstrated its utility in RLSs due to its portability, low cost, and lower reliance on traditional laboratory infrastructure. For example since the seminal ‘lab in a suitcase’ work, where on-site surveillance of Ebola mutation rates during the 2014-2016 outbreak in West Africa were performed using Oxford Nanopore’s MinION device [2], Nanopore sequencing has gone on to form a major component of Africa’s localized on-site COVID-19 surveillance response [3]. Similarly, the growing adoption of Nanopore sequencing for eDNA/taxonomic identification analysis in RLSs is evident from numerous recent examples, e.g., on-site taxonomic profiling of microbiota within the lava fields of the Canary Islands [4], in-field bio-surveillance subtyping of Venezuelan Equine Encephalitis virus in mosquitoes in Southern Florida [5], frog species identification in remote rainforests in Tanzania [6], and even proof-of-concept work to identify the previous locations of high-value military targets in simulated battlefields [7]. This growth in on-site, low-cost, simplified molecular analysis technologies has also contributed to discussions around the potential ‘democratization’ of eDNA testing outside of established scientific communities and infrastructure [8]. However, there still exist significant barriers for the uptake of eDNA nanopore sequencing as a ubiquitous tool in RLSs, due in part to the remaining high cost of upstream equipment for the collection, extraction, quantification, and processing of eDNA samples prior to their downstream analysis.

A major upstream requirement for eDNA Nanopore sequencing is the need to perform rigorous quantitation of eDNA samples by means of fluorometric methods [9]. Fluorometric quantitation relies on the use of fluorescent dyes that bind to Nucleic Acids, such as PicoGreen, whereupon their fluorescence increases thousand-fold times greater than their unbound state [10]. These techniques offer superior sensitivity and selectivity compared to traditional spectrographic methods [11] [12], making them a ‘best-practice’ tool for quantifying low-concentration DNA samples, which are often encountered in eDNA analysis pipelines. For example, during library preparation for Nanopore sequencing, quantitative analysis of nanogram masses of DNA must be made to ensure DNA molarity is within established ranges to maximize sequencing output [13]. Additionally, multiple fluorometry quality control (QC) checkpoints are present in most library preparation protocols to check for losses in yield throughout this expensive multi-stage process. Despite their simplicity and utility, the high cost of fluorescent readers is a significant barrier to performing eDNA sequencing in RLSs. For example, the predominant and popular Promega Quantus and Thermo Fisher Scientific Qubit retail for ∼$1500-3300 USD, respectively, making them a larger capital expenditure than ONT’s $1k USD MinION Nanopore sequencing device. The popularity of these fluorescent platforms also makes them susceptible to unexpected supply constraints, such as those experienced during the recent COVID-19 pandemic, where access to scientific equipment and consumables was a significant global challenge [14]. Therefore, portable, low cost, fluorometers, that can side-step centralized manufacturing bottlenecks, and be produced at the point-of-need, represent a major component to increasing access to eDNA sequencing in RLSs.

To address this need, we have developed the DIYNAFLUOR, a DIY plug-and-play nucleic acid fluorometer designed to offer a low-cost, portable, and easy-to-use solution for DNA quantification in RLSs. Built with readily available off-the-shelf components, simple 3D-printed parts, and a solder-free assembly process, The DIYNAFLUOR is designed to be manufactured, assembled, and validated within a day by non-technical users, enabling rapid, decentralized distribution in RLSs. Primarily designed to work with the established Qubit Broad Range (BR) and High Sensitivity (HS) DNA fluorometry kits, the sensitivity and accuracy of the DIYNAFLUOR to quantify nano and microgram per microliter levels of DNA is assessed herein. Laboratory verification was assessed by placement of a DIYNAFLUOR in a laboratory where comparative measurements to a commercial Qubit 4 fluorometer were made. To validate if assembly instructions were sufficiently clear, assessment of DIYNAFLUOR units assembled by researchers outside the preliminary design team was performed. To further address cost constraints in RLSs, we also investigated the capabilities of a custom, fluorescent assay to quantify eDNA extractions. This “extreme” low-cost assay uses the ubiquitous SYBR Safe DNA fluorescent dye at a per-measurement cost ∼10% that of commercial kits.

To highlight the DIYNAFLUOR’s applicability to RLS eDNA Nanopore sequencing, an experiment to extract eDNA from Australian lake water and to prepare and sequence metagenomic barcoding libraries targeting 18S and 16S ribosomal RNA genes was performed. All DNA quantification steps (i.e., sample purification and amplified NGS library quantification) were performed using the DIYNAFLUOR, followed by Nanopore sequencing. This approach successfully identified a variety of local Australian fauna, demonstrating the device’s effectiveness in supporting field-based genomic studies within RLSs. Finally, to demonstrate the DIYNAFLUOR’s extended use-cases, changes to the fluorescent configuration for RNA and Protein quantification are briefly discussed. By offering a cost-effective, portable, and reliable solution for fluorescent biomolecule quantification, the DIYNAFLUOR removes a major barrier to conducting molecular and genomic research in RLSs.

### Device Design

Figure 1(A) shows a DIYNAFLUOR unit. The main body of the device measures 130 × 71.5 × 44 mm. All structural parts are designed to be 3D printed on low-cost desktop 3D printers, without the need for supports and can be printed simultaneously on a 250×250 mm print bed. Assembly requires simple hand tools, with fasteners consisting primarily of machine screws. The DIYNAFLUOR has a maximum power consumption of ∼100 mW during measurements and can be powered and operated by USB connection to a computer, making it highly portable when paired with a laptop. Samples are measured by placing 0.5 mL thin-walled PCR tubes containing analyte in a Sample Well located on the top of the device. A separate Light Baffle can be placed over the tubes to prevent unwanted interference from outside light sources during fluorescent measurements.

**Figure 1.**
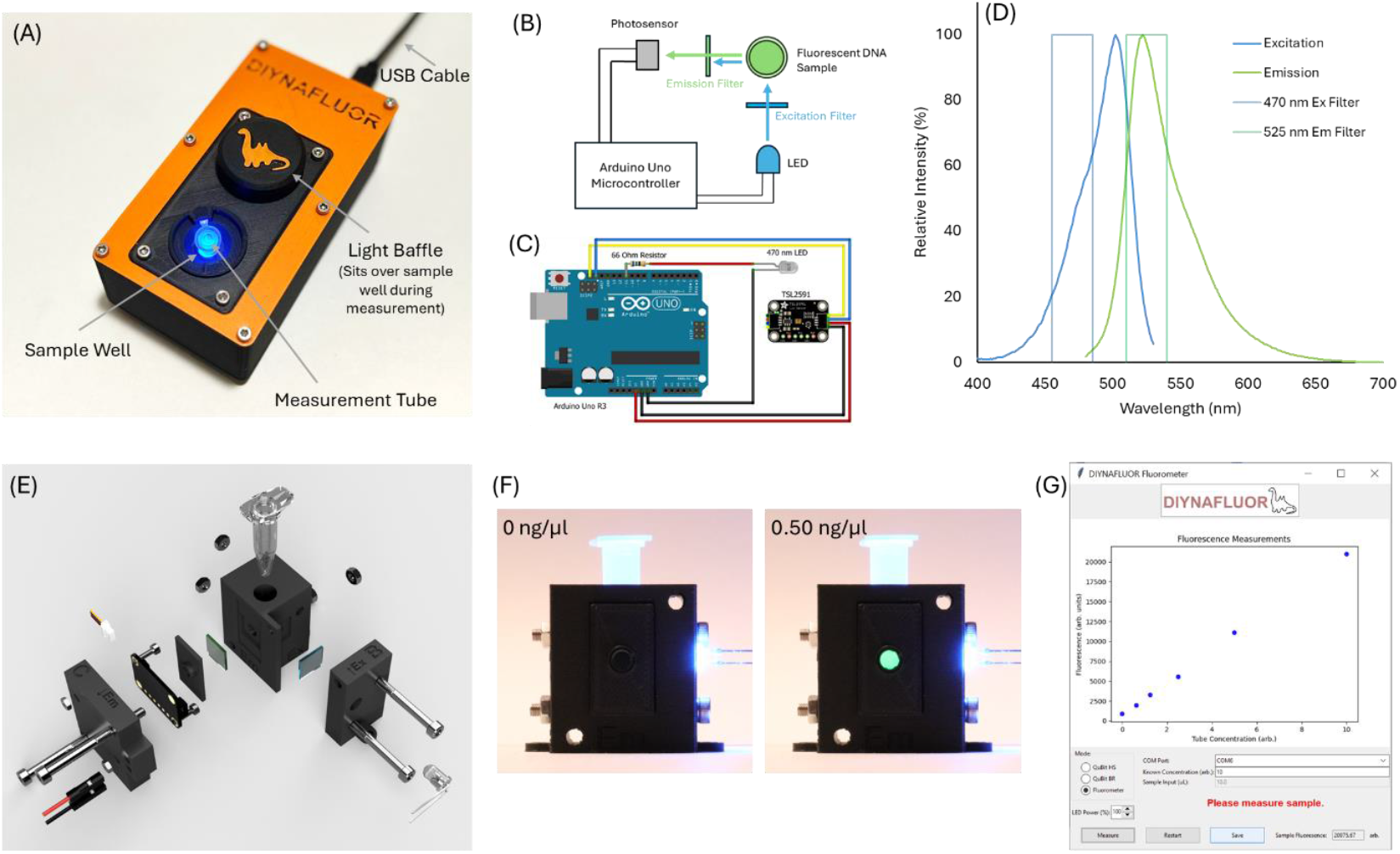
(A) The DIYNAFLUOR device. (B) A simplified systems architecture diagram. (C) Wiring diagram. (D) Emission and Excitation filters used in the DIYNAFLUOR overlayed on the fluorescent spectrum of Pico-green DNA dye. (E) An exploded 3D render of the “Filter Cube” that houses the core fluorescent detection components within the DIYNAFLUOR. (F) Images of the fluorescent signal produced within the filter cube for 0 and 0.50 ng/μL DNA samples using the Qubit HS reagents. (G) The DIYNAFLUOR control software.

Figure 1(B) shows a simplified schematic of the DIYNAFLUORs principle of operation, where a microcontroller activates the excitation LED and simultaneously takes a reading of the fluorescent emission signal by means of a photosensor. An Arduino clone (Uno R3) was chosen as the microcontroller for the DIYNAFLUOR due to its low cost, availability, existing library of code, and pre-soldered connection headers. Figure 1(C) shows a wiring schematic of the Arduino microcontroller circuit. A high power 470 nm, 12500 mcd, 5 mm round, through-hole LED, current limited with a 68 Ohm 5W series resistor, was connected to one of the PWM pins on the Arduino, allowing for variable excitation light intensities (at maximum power, the 68 Ohm resistor limits the maximum forward current to ∼28 mA to prevent damage to the Arduinos PWM pins). The LED and resistor were connected without soldering using 2.54 mm F-F and M-F DuPont connectors. Fluorescence measurements were performed with a Adafruit TSL2591 Digital Light Sensor, controlled by the Arduino over an I2C protocol. The TSL2591 sensor was chosen due to its low-cost, 600M:1 dynamic range, broad spectral response (400-1000 nm), programmable gain and integration times, Arduino library support and STEMMA QT connection port, which removed the need to solder connecting wires. All TSL2591 measurements performed in this manuscript were the average of 3x measurements made with gain set to ‘Max’ and an Integration Time of 600 ms.

Figure 1(D) shows the fluorescent excitation and emission spectrum of the PicoGreen dye used in the Qubit reagents with the excitation and emission bandpass filters used in the DIYNAFLUOR system. The excitation filter helps refine the excitation spectrum to 470 nm ± 15 nm, reducing the background signal of unwanted longer-wavelength green light from LED (the magnitude of which can vary slightly between LEDs). The 525 nm ± 15 nm emission filter prevents excitation light from reaching the TSL2591 sensor and is optimal for maximizing the fluorescent signal for the peak emission wavelengths of Pico-Green.

The optical configuration is constructed within a 3D printed Filter Cube, as shown as in the exploded diagram in Figure 1(E). Housed within the Main Body of the DIYNAFLUOR, the Filter Cube was configured with a 90° optical path to prevent direct illumination of the TSL2591 sensor by the excitation source, thereby minimizing background noise from any light able to pass through the emission filter. To further reduce unwanted signal, the Filter Cube was printed with matte black PLA filament to reduce background fluorescence that can occur in colored plastics, and to help diffuse reflected light across the surface of the TSL2591 sensor. The Filter Cube part containing the Sample Well also contains a light baffle that runs the length of the 470 nm LED to prevent side light leakage from reaching the TSL2591 sensor through joins between parts. The design aimed to position the 470 nm LED, TSL2591 sensor and optical bandpass filters as close as possible to the sample tube to reduce signal decay due to the inverse square law. A hole in the top of the Main Body sits flush with the top of the Filter Cube, allowing for access to the Sample Well. A “V-well” recession at the base of the sample well assists with consistent optical alignment of the 0.5 mL thin-walled PCR tubes. The LED and filters are modular and can be interchanged to support different fluorophores if needed. Figure 1(F) demonstrates the effectiveness of this filter cube design by showing the difference in fluorescent signal for a Qubit HS assay with 0 and 0.5 ng/μL DNA concentrations, as observed through the Emission Port with the TSL2591 sensor removed.

To simplify usability, a Graphical User Interface (GUI) software package was developed in Python, as shown in Figure 1(G). This custom software allows users to easily operate the DIYNAFLUOR device and provides basic data analysis and visualization of results. The GUI software, provided as an easy to install independent Windows 10/11 package, Arduino microcontroller code and 3D print files are available on our GitHub Repository [15], as well as a detailed Bill of Materials (BoM), and Build Instructions Manual – containing 3D printing guidance, step-by-step assembly instructions and a qualification assay.

## Results and Discussion

### Assessment of Fluorescent Response and Dynamic Range

To assess the fluorescent response to DNA at different concentration and to determine the dynamic range of the DIYNAFLUOR system, triplicate measurements were performed for serial dilutions of DNA for the Qubit HS and BR kits across a range of 0-0.5 and 0-5 ng/μL, respectively. These concentrations fall within the working ranges of these assays as defined by Thermo Fisher Scientific. It must be noted that all concentrations mentioned in this, and the following “Sensitivity” section, refer to the ‘in-assay’measurement tube concentrations, which is the final concentration of 10 µL of stock DNA from the dilution series added to 190 µL of the Qubit master-mix. Values can be converted to the stock dilution series concentrations by multiplying the in-assay concentration values by 20. Figure 2 shows raw fluorescent data for the 2-fold dilution series of the Qubit HS and BR reagents measured by both DIYNAFLUOR and a commercial Qubit 4 fluorometer. A linear regression for both the HS and BR assays measured by DIYNAFLUOR provide an R^2^ of greater than 0.999, indicating a highly linear fluorescence response to DNA input concentrations across this working range of DNA input. Similar linearity was seen in the commercial Qubit platform, which achieved an R^2^ values greater than 0.9989 for both assays. A quarter-fold dilution series of DNA for both kits showed similar results and are provided in Fig S1. Importantly, the dynamic range (the difference in fluorescent signal between the blank and highest DNA concentration) for the HS assay as measured by DIYNAFLUOR was ∼23,550 units, equivalent to ∼36% of the available dynamic range of the 16-bit TSL2591 optical sensor. Assuming a perfectly linear fluorescence response to DNA concentration, and ignoring random measurement error, this dynamic range provides a theoretical maximum resolution of the DIYNAFLUOR of ∼21 fg/µL.

**Figure 2.**
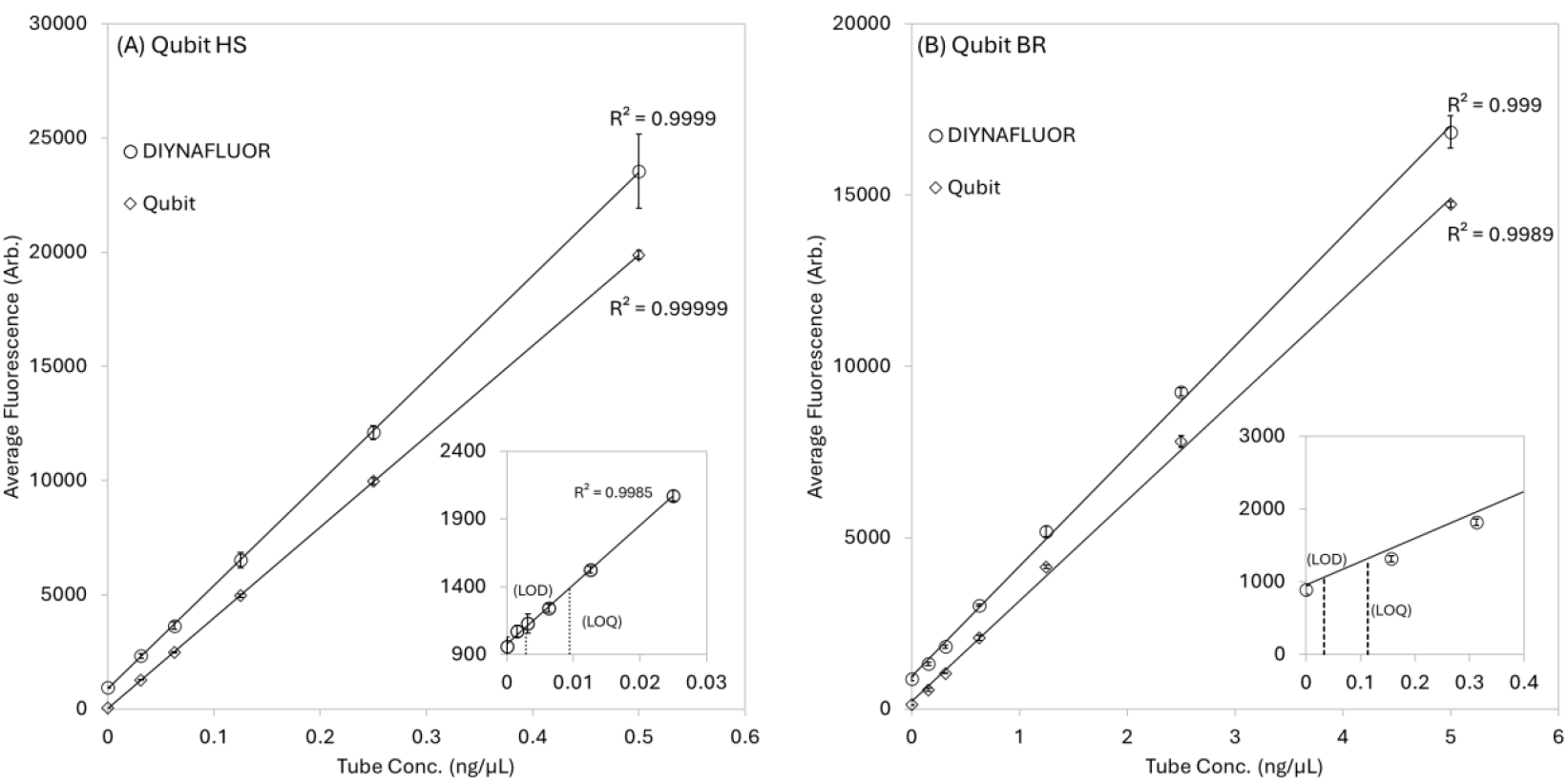
Assessment of the linearity and sensitivity of the Qubit HS and BR reagents when measured by DIYNAFLUOR and Qubit 4. Calculated LOD and LOQ values are highlighted by dashed lines. (A) Raw fluorescent data of 3n replicates of a 2-fold dilutions series of DNA using the Qubit HS assay reagents. The inset graph shows 3n replicates of a 2-fold dilutions series of DNA across the low range of the assay (B) Raw fluorescent data of 3n replicates of a 2-fold dilutions series of DNA using the Qubit BR assay reagents. The inset graph shows a zoomed section of the low range of the assay.

### Sensitivity

We define sensitivity as the empirically determined Limit of Detection (LoD) and Limit of Quantitation (LoQ), which are the lowest concentration that can be measured with statistical significance above baseline noise, and the lowest concentration of a substance that can be reliably measured and quantified with a known degree of accuracy and precision, respectively. As described by the Eurachem Method Validation Working Group [16], the LOD and LOQ for a measurement platform should be obtained from the standard deviation, σ, of 10 replicate blank measurements, where 3.3σ, is the LoD and 10σ is the LoQ. For example, the LoD and LoQ for the Qubit HS assay was measured to be 128, and 426 arbitrary fluorescence units, respectively. Blank measurements for all DIYNAFLUOR systems presented in this study consistently fell within ∼500-1500 units for the HS assay, as such we don’t expect the baseline variance to change significantly across units, but this can be easily assessed if required by the end user. Assuming a perfectly linear fluorescence response to DNA concentration, the in-assay LoD and LoQ can easily be converted to units of concentration by substituting the arbitrary fluorescence values into the slope of the average minimum and maximum fluorescence values obtained from the dilution series from Figure 2. The inset graphs in Figure 2(A) and (B) show the LoD and LoQ values at the lower range of the assays, which were 0.003 and 0.009 ng/μL, respectively, for the HS assay and 0.034 and 0.113 ng/μL, respectively, for the lower sensitivity BR assay.

The same 10x replicate blank measurements were performed on a Qubit 4 for comparison. The LoD and LoQ for the HS and BR assays were measured as 0.0002 and 0.0007, and 0.002 and 0.007 ng/μL, respectively, when using the slope obtained from the 2-fold dilution series measurements.

This ∼13-16x increase in sensitivity compared to the DIYNAFLUOR is likely due to the higher quality optics and more consistent tube placement afforded by the commercial grade manufacturing processes used in the Qubit system, which are more readily able to handle the small optical aberrations in the 0.5 ml thin-walled PCR tubes, and small deviations in tube placement, compared to the DIYNAFLUOR’s 3D printed and off-the-shelf parts. These variations can act to change the optical pathlength of the fluorescent signal between measurements. Fig S10 provides further analysis of this source of random error, finding a maximum random error in the fluorescent signal of ∼4.5% across the Qubit HS working range. As will be demonstrated, despite this error, the DIYNAFLUOR’S sensitivity and accuracy is adequate for most molecular biology processes, such as QC-ing sequencing library preparations.

### 2-Point Calibration Accuracy (Bias and Precision)

As outlined earlier, the Qubit HS and BR assays produce a highly linear fluorescent response across the working input DNA concentrations. We can therefore use a simple linear 2-point calibration methodology to determine unknown DNA concentrations as follows.

Using two fluorescent standards, S_1_ and S_2_, where S_1_ is a blank and S_2_ is a DNA standard of known concentration, the unknown DNA concentration, C, is calculated by

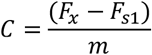

where F_x_ is the fluorescent value of the unknown DNA sample, F_s1_ is the fluorescent value for the blank standard, S_1_, and m is the gradient obtained from a two-point calibration. m can be acquired from the measurements of the two standards by

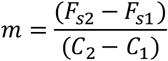

where F_s2_ is the fluorescent value of S_2_, and C_2_ and C_1_ are the DNA concentrations of S_2_ and S_1_, respectively.

For the HS for the BR Qubit assays, these values are 0.5 and 5 ng/μL for the S_2_ standards, respectively, and 0 ng/µL for both S_1_ standards.

The Qubit reagents allow for 1-20 µL of DNA to be added to the fluorescent master mix to a total volume of 200 µL to allow for measurement of sample DNA concentrations that may fall above or below the working range of the fluorometer. Therefore, to calculate the sample concentration of DNA, C_s_, the dilution factor, df, must be accounted for, where

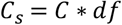

df is easily calculated by dividing the total reaction volume (200 µL), by the input DNA volume.

The DIYNAFLUOR software incorporates a protocol that guides the user through the 2-point linear calibration followed by measurement of their samples. Briefly, either the BR or HS kit is selected, (C_2_ concentration values for either kit will be automatically applied to the analysis) and measurement of the S_1_ and S_2_ standards are then performed. The input of the unknown samples df can be input prior to the measurement of the unknown DNA sample to provide its stock concentration.

To assess the accuracy of this 2-point calibration methodology, the DIYNAFLUOR’s absolute bias was measured. Eurachem [16] describes the absolute bias of an analytical platform as the difference between the mean of the measured results and the value of the true sample being measured. A suggested minimum of 10 samples should be tested to ascertain bias in an analytical platform. We report bias as the average absolute bias for 3n replicates over a two-fold dilution series of measurements across the working ranges of the BR and HS assays (7 concentrations were measured from 100-0 ng/μL for the BR assay, and 6 concentrations were measured from 10-0 ng/μL for the HS assay). This global approach allows for variations in bias across the working range of the assay to be accounted for. In simplistic terms, the bias values provided here represent the average expected error in any measurement made by DIYNAFLUOR for the BR and HS assays. Similarly, precision is defined as the average standard deviation for the 3n replicates at each concentration over the dilution series and is presented within brackets after the reported bias values.

Figure 3 shows the measured DNA concentrations using a 2-point calibration methodology for both the HS and BR Kits. Comparative measurements for the same samples were also performed on a Qubit 4.0 device to assess the performance of both fluorometers. A highly linear trend for both assays and measurement platforms was observed with the bias and precision for the HS and BR kits being 0.018 (0.093) and 0.050 (0.549) ng/μL, for the DIYNAFLUOR measurements, respectively, and -0.008 (0.044) and -1.424 (0.353) ng/μL, for the Qubit HS and BR assays, respectively. The larger bias values in the BR kits were expected due to the larger dynamic range of DNA concentrations across which this assay operates. It must be noted that the commercial Qubit fluorometer utilizes a proprietary 2-point calibration methodology based on a Hill equation, which can account for small deviations from linearity in the fluorescent response to analyte concentrations across the range of Qubit fluorescent assays [17]. This variation is evidently small for the HS assay reagents, as we could not identify any significant deviation from linearity across the dilution series within the DIYNAFLUOR data that the Qubit’s Hill equation claims to account for. However, the small deviation from linearity seen in the 25 and 50 ng/μL measurements in the DIYNAFLUOR’s BR dilution series, which appears to be handled better in the Qubit measurements, are evidence of some non-linearity in that assay. Regardless, this deviation represents an error of less than 3 ng/μL. Insets in Figure 3 show zoomed data at the lower working concentration rage, highlighting the LoD and LoQ in terms of measurable DNA concentration from stock solutions based on the previously established sensitivity values, which were 0.028 and 0.094 ng/μL for the HS assay and 0.34 and 1.13 ng/μL and BR assay, respectively. Additional concentration measurements across a quarter-fold dilution series are shown in Fig S2.

**Figure 3.**
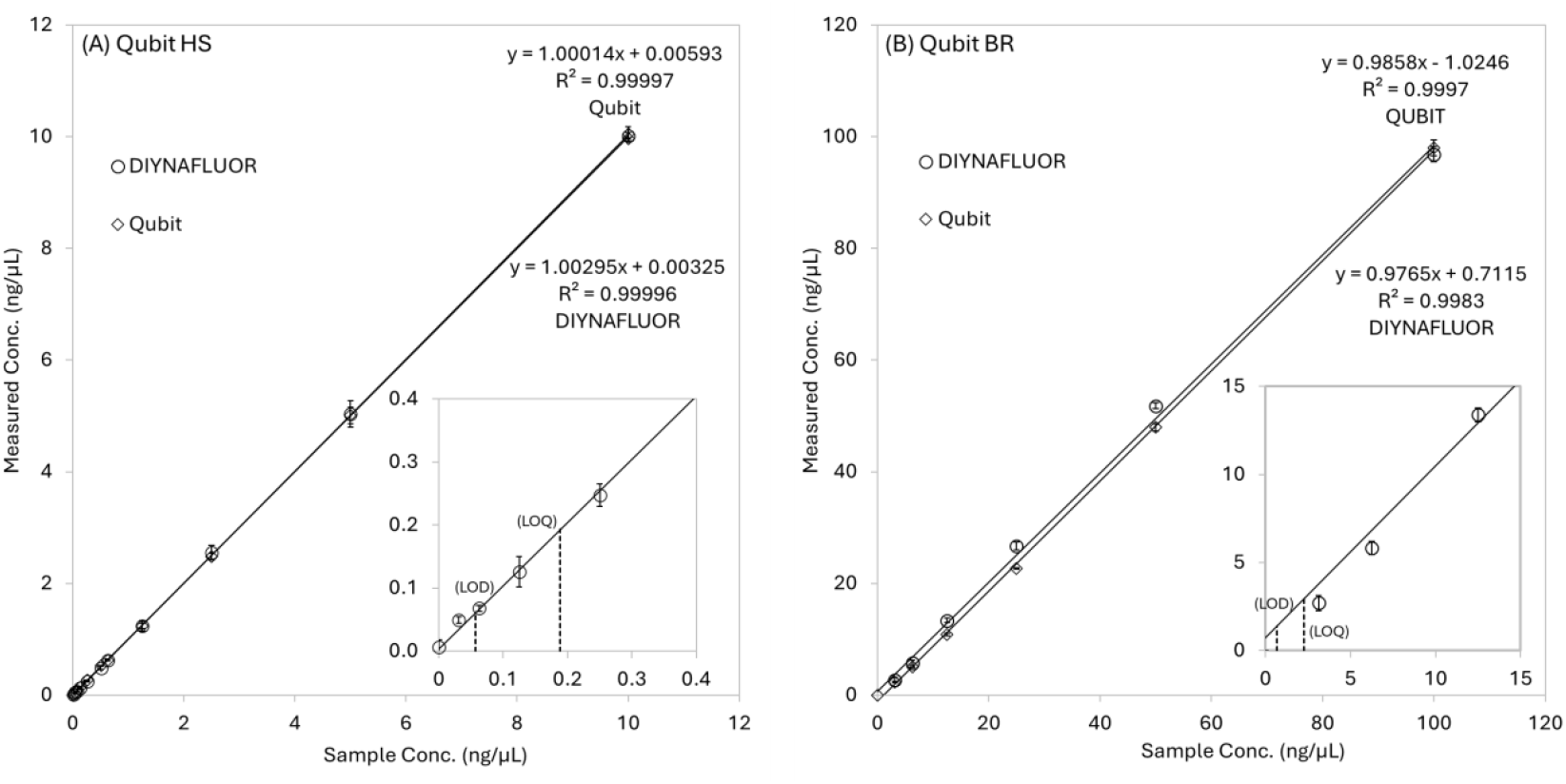
DIYNAFLUOR and Qubit 4 measurements of DNA samples with known concentrations (same samples used in Figures 1 and 2) using a 2-point standard curve calibration procedure. (A) A high correlation was observed between the two techniques using the HS Qubit reagents, with the data points, error bars and linear regressions for both techniques being almost indistinguishable. (B) Deviation between the two techniques for the BR Qubit reagents was broader than the HS reagents, however both techniques deviated from the known concentration to a near equal extent suggesting the variance is inherent to the assay. Insets in both graphs show low range data for both assays with the LoD and LoQ shown as dashed lines.

In real-world DNA sample extractions, small amounts of impurities in the form of extraction chemicals or unwanted biological source material are often present in the final purified product. To assess the DIYNAFLUOR’s accuracy in a “real-world” setting, and to ascertain if impurities affect the accuracy of the Qubit HS assay, small amounts of common DNA extraction chemicals and biological materials, (e.g., Ethanol and Protein (BSA)), were spiked into Qubit HS DNA assays. Fig S3 shows measurements performed by DIYNAFLUOR and Qubit 4, finding similar deviations between the two technologies. Interestingly, in some cases the impurities were found to cause >20% deviation from the expected value, which we have previously not seen reported in the literature. This observation reinforces the need to optimize DNA extraction protocols to ensure purity and accurate quantification for downstream analysis.

### Laboratory Verification

To verify the DIYNAFLUOR was able to perform as intended in a research environment, a unit was placed in a biotechnology laboratory within The University of Queensland for ∼2 months. Research staff and students working in the laboratory and who were familiar with DNA fluorometry techniques were asked to perform a comparative measurement on the DIYNAFLUOR any time they performed a Qubit DNA HS assay during their regular laboratory work. Users were trained on the device, and asked to follow the Qubit HS DNA protocol, as defined by Thermo Fisher Scientific, to prepare their samples for analysis. Calibration and samples tubes were measured first by Qubit 4, and then immediately on the DIYNAFLUOR unit. Only the date, time, input volumes and measured concentrations were recorded. Over the verification study, 209 samples were measured. Figure 4(A) shows the comparative data obtained from the verification study, with the inset showing zoomed data for the 0-10 ng/μL regime, where most measurements fell. Linear regression of comparative measurements gave an R^2^ of 0.9963 and gradient of 0.9447, indicating a highly linear correlation between the two platforms. Measurements that reported as being greater than the maximum working range of the assay by Qubit analysis were excluded, and those that reported below the minimum working range by Qubit analysis were reported as 0 ng/μL for the Qubit result.

**Figure 4.**
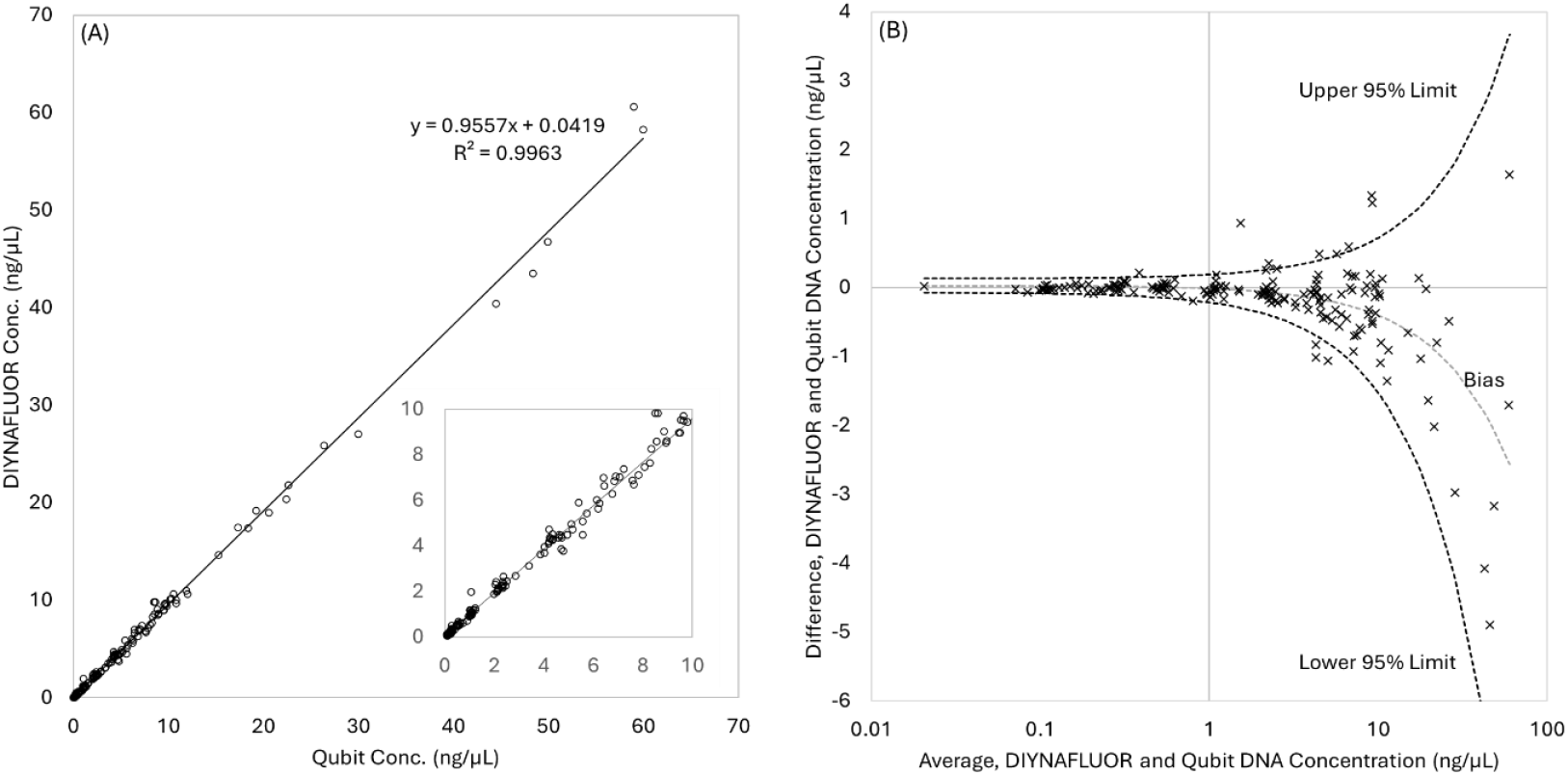
Verification data collected during the 2-month placement of a DIYNAFLUOR in a biotechnology laboratory. (A) Shows same-tube comparison of HS Qubit DNA assays performed by both DIYNAFLUOR and Qubit. The inset graph shows a zoomed section of the low concertation samples. (B) A Bland-Altman plot showing the 95% Upper and Lower Confidence of the variation between the two measurement platforms.

To compare the variance in the two techniques, a Bland-Altman plot was generated. A Bland-Altman plot assesses the agreement between two measurement techniques by plotting the differences against the averages of multiple paired measurements to identify bias, variability, and any trends or outliers. To account for the greater variance in the data at larger values, a modified Bland-Altman analysis that accounts for relationships between the variance and magnitude of the measurement was performed [18]. Figure 4(B) shows the Bland Altman plot, with 95% Upper and Lower Confidence Limits showing the expected variation of the two techniques across a log concentration range of 0.01 – 100 ng/μL. Of note is the small variance at the lower concentration measurements, where values below 1 ng/μL are only expected to deviate by ∼±0.2 ng/ul between the two fluorometers.

We must report that early in the verification study, users reported significant deviations between the Qubit and DIYNAFLUOR measurements. It was quickly established that this was the result of different geometries in a newly purchased lot of 0.5 mL thin-walled PCR tubes provided by Thermo Fisher Scientific. While the different tube geometries did not affect the Qubit platform measurements, mixing tube geometries would lead to significant deviations when used interchangeable in the DIYNAFLUOR platform, i.e., the same DNA concentration measured within different tube geometries would give significantly different raw fluorescent values. It was quickly realized that this was due to the slightly different fluorescent pathlengths between the two geometries and was easily remedied by ensuring that only one type of tube was used for this study. Fig S4 shows the difference in these tube geometries, which we have identified as being interchangeably distributed on a lot-to-lot basis by Thermo Fisher Scientific. We therefore recommend that 0.5 mL thin-walled PCR tubes are checked for variations in tube geometry, or that a singular supplier of 0.5 mL thin wall PCR tubes, such as those produced by Axygen, are used for DIYNAFLUOR measurements. Early measurements where mixed tube geometries were identified as being the cause of the aberrant measurements were removed from the analysis.

### Build Instruction Validation and Qualification

A Build Instruction Manual was produced to help end users with assembly and qualification of a DIYNAFLUOR device. This manual provides guidance for desktop 3D printing of components, visual step-by-step assembly instructions, software installation and a Qubit-based qualification assay. To validate the clarity of the Build Instruction Manual, three volunteer researchers with backgrounds in using fluorometric techniques, but a self-defined limited backgrounds in electronic device design, were provided with kits containing the individual, unassembled component (including 3D printed parts), assembly tools, software, the Qubit reagents, laboratory tools to perform the qualification assay, and the Build Instruction Manual. The volunteer researchers were asked to follow the Build Instruction Manual and report the results of the qualification assay, which was a non-replicate, 2-fold serial dilution using the Qubit HS DNA assay (the same assay as outlined in the “Assessment of Fluorescent Response and Dynamic Range” section). Figure 5 shows the qualification assay data for 6 DIYNAFLUOR units, 3 built and tested by the primary design team, and 3 built and tested by the volunteer researchers. One of the design team units had its 3D printed parts produced on a Bambu Lab A1 printer to compare performance with the other units, which had parts printed on a Creality Ender 3. All units produced similar linear fluorescent responses across the DNA dilution series, with Dynamic Ranges of ∼25,000 units and R^2^ values greater than 0.996. Variations between units are most likely due to semiconductor process variation in the LED output intensities and the sensitivities of the TSL2591 sensors. The volunteer researchers were all able to assemble a DIYNAFLUOR and perform the qualification assay within 1 day. Following the build, feedback was taken from the volunteer researchers to improve any concerns with the clarity of the Build Instruction Manual, which is provided at [15].

**Figure 5.**
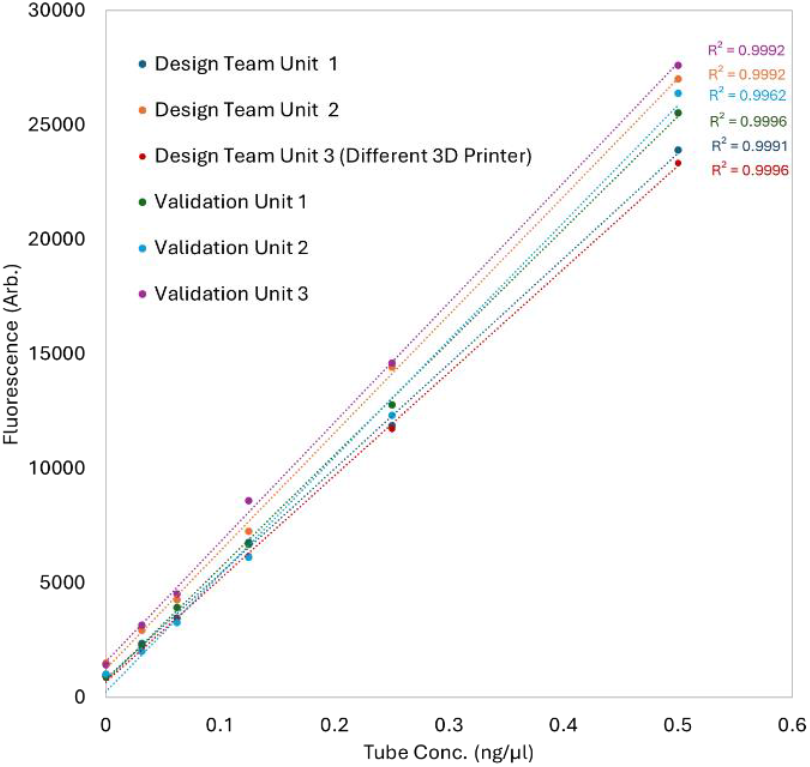
A 2-fold dilution assays using the HS Qubit reagents used to qualify six DIYNAFLUOR devices. Three devices were assembled and qualified by the lead designer and the other three were assembled and qualified by three scientists outside the design team following a build guide document.

### Extreme Low Cost SYBR Safe Assay for eDNA Extractions

To further help reduce barriers to eDNA analysis in RLSs, we developed and assessed an “extreme” low cost fluorometric assay that uses SYBR Safe. SYBR Safe is a low cost and readily available fluorescent NA dyes used ubiquitously to visualize DNA in electrophoretic gels. It is also claimed as a safer alternative to other low-cost DNA binding fluorogenic dyes [19]. To assess SYBR Safe for DNA quantitation, fluorescent readings across a 2-fold dilution series from 0–5 ng/μL were performed, as shown in Figure 6(A). The linear working regime of the assay was confirmed between 0-0.5 ng/μL, enabling a 2-point calibration methodology with the DIYNAFLUOR software, as described earlier. The bias was found to be -0.128 (0.241) ng/μL across the working regime. Using the previously described sensitivity assessment methodology for the Qubit assays, the in-assay LoD and LoQ were found to be 0.014 and 0.048 ng/μL, respectively. SYBR Safe is also known to fluoresce when bound to RNA, although at a much lower efficiency. Assessment of assay robustness to RNA was performed by spiking equal concentrations of RNA into DNA samples across a range of concentration, finding a negligible increase in signal when compared to a DNA only control (See Fig S5). For users concerned about RNA signal skewing results, RNAse treatment prior to measurement should be considered. While this assay is less sensitive and accurate than the Qubit HS assay, it is costed at only ∼$0.13 USD per assay. This is approximately 1/10th the cost of commercial NA assays, at ∼$1 USD per-assay (see [15] for a BoM and costings). We suspect this low-cost, custom assay may be an attractive alternative for STEM educators, or LRSs in times of extreme resources or supply constraints.

**Figure 6.**
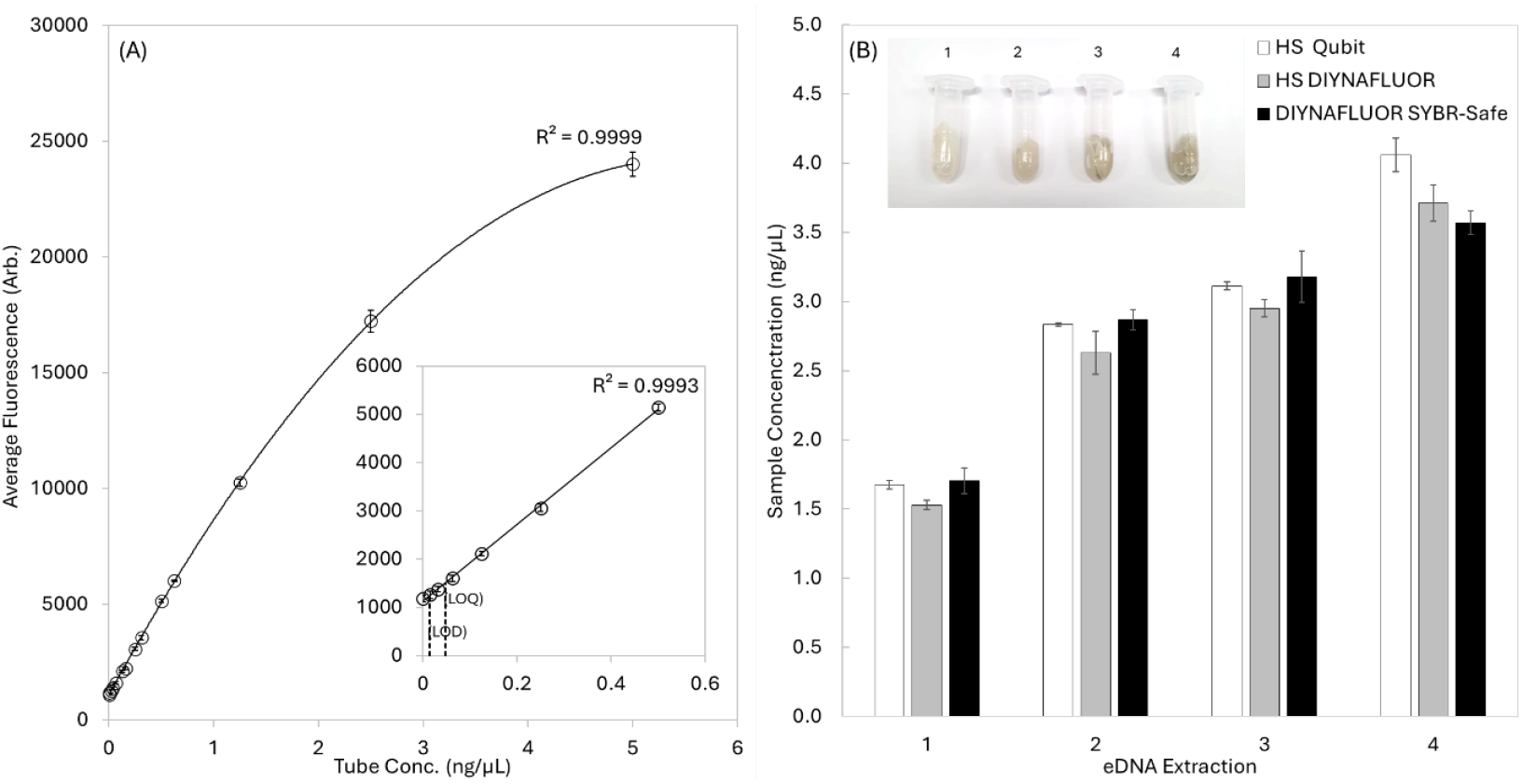
(A) A 2-fold dilution series of lambda-phage DNA measured by DIYNAFLUOR. A non-linear regression to a 2^nd^ order polynomial was used to fit the data. The inset graph shows the zoomed in, linear regime of the assay, and a corresponding linear fit. (B) Comparisons of measured concentrations of eDNA extracted from four samples of UQ lake water using HS Qubit reagents by both Qubit and DIYNAFLUOR, and by 2-point standard curve analysis within the linear regime of the SYBR Safe DIYNAFLUOR assay. The inset image shows the four 0.45 μm PVDF filters used to concentrate the lake water eDNA during the ATL buffer treatment stage of the extraction protocol.

To assess the utility of the assay for eDNA quantitation, eDNA purified from a local freshwater lake was quantified with the low cost SYBR Safe assay and compared to DIYNAFLUOR and Qubit 4 measurements using the Qubit HS reagents. Measurements were performed on 3n replicates of 10 µL of each sample. The same tubes used to measure the Qubit HS Assay were measured in both the Qubit 4 and DIYNAFLUOR. Figure 6(B) shows the results of the four samples measured across the three techniques, with the largest variance seen in sample four, between the Qubit HS measurement and the DIYNAFLUOR SYBR Safe measurement, which had an absolute difference of 0.49 ng/μL, representing a 12% variance between techniques. For eDNA QC applications, this variance is likely acceptable for most applications and the overall good agreement for all methods demonstrates the real-world application of this extreme lost cost assay.

### Nanopore eDNA sequencing and Identification of local Australian fauna

To assess the ability of the DIYNAFLUOR to QC eDNA library preparations, comparative DIYNAFLUOR/Qubit measurements of the QC steps within a Nanopore Native Barcoding Library protocol were performed on 16S and 18S metagenomic libraries generated from the four eDNA lake extractions described earlier. The library preparation consists of four QC fluorometry steps at amplicon generation, barcoding, adapter ligation and final pooled library and is outlined in the material and methods section. Table 1 shows the comparative measurements at the different QC stages of the library preparation. In general, all measurements agreed well, with either the Qubit or DIYNAFLUOR values being appropriate for QC calls or molarity calculations throughout the assay. With amplicon libraries generated, a Nanopore sequencing run was performed as described in the materials and methods and the following bioinformatic protocol was employed.

**Table 1.** QC data obtained by DIYNAFLUOR and Qubit during the library preparation for eDNA nanopore sequencing and Australian fauna identified from the metabarcode sequencing results.

| Nanopore Library Prep QC Measurements |  |  |  |  |  |
| --- | --- | --- | --- | --- | --- |
| QC Stage |  | Sample | Input Volume (uL) | Qubit (ng/uL) | DIYNAFLUOR (ng/uL) |
| Amplicon Library PCR |  | Extraction 1 16S | 2 | 35.4 | 33.41 |
|  |  | Extraction 2 16S | 2 | 35.2 | 32.68 |
|  |  | Extraction 3 16S | 2 | 34.8 | 34.76 |
|  |  | Extraction 4 16S | 2 | 32.8 | 31.67 |
|  |  | 16S NTC-1 | 2 | Too Low | 0.20 |
|  |  | 16S NTC-2 | 2 | Too Low | -0.13 |
|  |  | Extraction 1 18S | 2 | 25.6 | 23.18 |
|  |  | Extraction 2 18S | 2 | 24.4 | 22.89 |
|  |  | Extraction 3 18S | 2 | 17.1 | 15.90 |
|  |  | Extraction 4 18S | 2 | 16.8 | 14.34 |
|  |  | 18S NTC-1 | 2 | 0.185 | 0.25 |
|  |  | 18S NTC-2 | 2 | 0.278 | 0.60 |
| Barcode Ligation End-Prep (16S and 18S Amplicons for each extraction pooled) |  | Extraction 1 | 1 | 6.20 | 6.20 |
|  |  | Extraction 2 | 1 | 6.30 | 5.91 |
|  |  | Extraction 3 | 1 | 5.68 | 4.91 |
|  |  | Extraction 4 | 1 | 6.34 | 5.22 |
| Bead Cleaned Pooled Barcoded Library |  | Total Pooled Library | 1 | 6.30 | 5.80 |
| Bead Cleaned Nanopore Sequencing Adapter Ligation |  | Total Pooled Library | 1 | 9.10 | 8.43 |
| Identified Australian Fauna |  |  |  |  |  |
| Number of Nanopore Reads |  |  |  |  |  |
| Extraction 1 | Extraction 2 | Extraction 3 | Extraction 4 | Scientific Name | Common Name |
| 3 | 10952 | 13503 | 20177 | Rhinella marina | Cane toad |
| 147 | 510 | 91 | 39 | Trichosurus vulpecula | Common brushtail possum |
| 113 | 32 | 174 | 51 | Anguilla reinhardtii | Speckled longfin eel |

Metagenomic libraries created using 16S and 18S rRNA gene sequences are inherently complex due to the high diversity and degeneracy of these sequences, which often include multiple variants with varying levels of sequence homology. This complexity can make it challenging to easily identify and quantify species present in environmental samples and creates an informatic bottleneck by analyzing the same read sequence multiple times. Therefore, an effective bioinformatics strategy is necessary to reduce the sequence space while still retaining the ability to distinguish between different species. Such a strategy must balance the need for specificity with the practical limitations of sequence analysis tools and computational resources.

To address this, a bioinformatics pipeline was employed to preprocess and analyze the FASTQ reads from the metagenomic libraries. Initially, FASTQ files from all barcoded samples were merged, and reads outside the desired size range (less than 100 bp or greater than 500 bp) were removed to exclude potential artifacts and off-target sequences. Overrepresented sequences were identified using ‘fastx_collapser’, which clustered identical reads to streamline the dataset. Further refinement was achieved through custom Python scripts that removed sequences with high similarity, defined as sequences differing by three or fewer mismatches or indels of up to two base pairs. Homology assessments were conducted by self-aligning the read clusters using BLAT, which allowed for the identification and exclusion of redundant sequences. The filtered read clusters were then indexed using Bowtie2, and the original reads were mapped to these clusters to facilitate coverage analysis with Bedtools’ ‘genomecov’ function. This approach enabled the estimation of species abundance by analyzing the proportion of 16S and 18S reads mapping to each cluster. Species identification was completed using NCBI nucleotide BLAST, providing a detailed taxonomic profile of the samples.

Manual evaluation of the 45 read clusters that passed the filtering criteria revealed a diverse array of local Australian fauna and organisms commonly found in or near pond environments. Notable identifications included the Australian Common brushtail possum, Cane toad, and Speckled longfin eel, alongside other expected freshwater species (See Table S1 for all identified species). Table 1 shows the number of reads for each sample that aligned to these notable Fauna, showing generally similar reads for each species across each extraction. The significant variance in the Brushtail possum reads for sample 1 are most likely due to the generation of longer amplicons in that sample skewing the molarity calculations (see Fig S7 for a gel of the Amplicon products). Overall, quantification and equimolar pooling of the four amplified and barcoded libraries base on DIYNAFLUOR measurements ensured that the four libraries were sequenced in equivalent proportions (Table S2 shows all barcoded samples were read ∼200k each), highlighting the DIYNAFLUORs’ capability for accurate and reliable quantification as a cost-effective and portable solution for eDNA analysis in RLSs.

### Other Assays

Finally, to demonstrate the broader utility of the DIYNAFLUOR for general molecular biology use, modifications were made to two units’ optical configurations to enable measurements with the Qubit RNA Broad Range and Protein Assays. This was achieved by simply changing the LED and bandpass filters to suit fluorophores used in these assays. For example, the Qubit RNA Broad Range assay uses a proprietary red fluorophore that is excited by 600-645 nm light and emits at 665 – 720 nm. As shown in Figure 7(A), by configuring a DIYNAFLUOR to use a 624 nm LED, and narrow bandpass filters at 630 nm for excitation and 682.5 nm for emission, measurements of a 2-fold dilution series across a range of 0-5 ng/μL of RNA were able to be made. Similarly, a configuration for the Qubit Protein Assay was able to measure protein across a 2-fold dilution series from 0-20 ng/μL, as shown in Figure 7(B), using the same 470 nm LED and 470 nm bandpass excitation filter as the Qubit DNA setup, but changing the emission filter to a 630 nm bandpass filter. This configuration met the spectral requirements of the proprietary fluorophore used in that assay, as outlined by Thermo Fisher Scientific. The BoM, available at [15] contains the optical components used for these configurations. Data for both configurations was non-linear least-squares fit against a modified Hill Equation using the Solver plugin in Excel [20], giving R^2^ values > than 0.9999 for each assay. While these non-linear assays are not supported by the DIYNAFLUOR software, there is an option to use the system in a “Fluorometer” mode, which allows for raw fluorescence values to be recorded for manual post-processing. Fig S9 shows the use of the Fluorometer mode to measure fluorescence for other assays using the standard ‘DNA’ DIYNAFLUOR configuration, including RNA with the low-cost SYBR Safe assay, Fluorescein isothiocyanate (FITC), and Promega’s commercial Quantifluor ONE dsDNA system. This versatility and adaptability of the DIYNAFLUOR system for other common fluorescent quantifications assays demonstrates its utility for wider use in RLS research environments.

**Figure 7.**
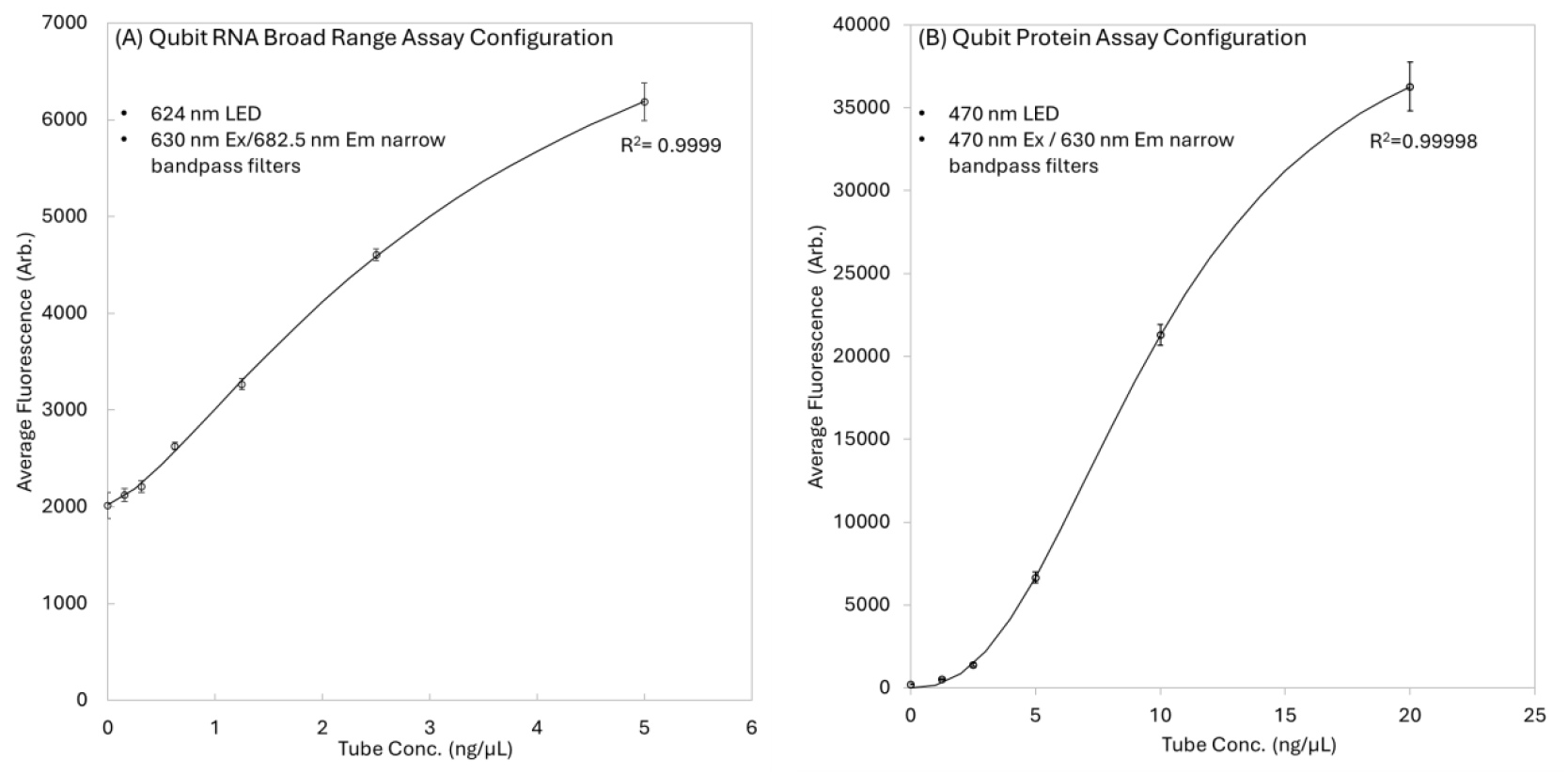
Different optical configuration of the DIYNAFLUOR. (A) A 2-fold RNA dilution series using the Qubit BR RNA reagents in a modified DIYNAFLUOR configuration targeting 630ex/682.5em fluorescent signals. (B) A 2-fold protein dilution series using the Protein Qubit Protein Assay reagents in a modified DIYNAFLUOR configuration targeting 470ex/630em fluorescent signals.

## Material and Methods

### Modelling and 3D Printing

All 3D printed components were designed and modelled in Fusion 360 (Autodesk). Design team units 1 & 2, and Validation Units 1 through 3 were printed on a Creality Ender-3 (Creality) with a 0.4 µm hot end. Design team unit 3 was printed on a Bambu Lab A1 (Bambu Labs) with 0.4 µm hot end. Simplify3D 4.1.2 (Simplify3D) software was used to slice the models for the Creality Ender-3 prints and Bambu Studio 1.9.7.2 (Bambu Lab) was used for the Bambu Lab A1 (Bambu Lab) prints. 3D parts were sliced for printing using near-default settings for the “medium” or “standard” settings in Simplify 3D or Bambu Studio, respectively. In general, prints were made with layers at 200 µm and without supports. Due to the differences in 3D printers and filaments, exact slicer settings are not provided, but general guidance for 3D printed parts is provided in the Build Instruction Manual available at [15]. All Creality Ender-3 and Bambu Lab A1 3D prints were made with PLA Matte Black (3D Fillies), and PLA Matte Black (Bambu Lab), respectively, excluding the non-functional dinosaur logo and colored “vanity plate” parts.

### Microcontroller and GUI Software

Arduino microcontroller code was written, compiled and uploaded with Arduino IDE Version 2.3.2. The GUI software was written in Python 3.12. The standalone Windows 10/11 package of the GUI was generated with PyInstaller 6.5.0. Source code and the standalone GUI software can be found on our GitHub Repository [15].

### Qubit Assays

Qubit quantification kits (Thermo Fisher Scientific) used in this study were as follows: Qubit 1X dsDNA HS Assay Kit, Qubit dsDNA BR Assay Kit, Qubit RNA BR Assay Kit, Qubit Protein BR Assay Kit. The manufacturers’protocols were followed relevant for the Qubit kit in use. The S2 standard were used for all DNA and RNA assays, being diluted into LTE buffer (10 mM TRIS (Sigma Alderich), 0.1 mM EDTA (Sigma Alderich), pH 8.0), or the relevant S1 standard, for all dilution series shown. In all assays (except for the Laboratory Validation study) 10 µL was added to 190 µL of the prepared Qubit master mixes. Similarly, the S3 standard was used in the same fashion for the Qubit Protein Assay experiments.

### Low Cost SYBR Safe Assays

The low-cost SYBR Safe Assay followed a protocol identical to the commercial Qubit dsDNA BR Assay. In brief, a SYBR Safe dye stock was made by diluting 10000x SYBR Safe (Thermo Fisher Scientific) to 200x in LTE Buffer. For each sample (including the 2 standards), 1 µL of stock dye was added to 199 µL of LTE buffer to make a master-mix. 190 µL of the master-mix was added to 0.5 mL thin-walled PCR Tubes (Axygen) and to this 10 µL of DNA solution added, followed by vortex mixing, bench-top centrifugation, and incubation for 1 min at room-temperature prior to measurement. A two-point calibration was performed by measuring 10 µL of two standards, being LTE and a 10 ng/μL solution of Lambda-Phage DNA (Thermo Fisher Scientific) in LTE, respectively.

### eDNA sampling

eDNA was collected and purified following guidance from [21]. Briefly, 500 mL of lake water was collected from The University of Queensland, St Lucia Campus lakes (27.500394368815467° south, 153.01496914473674° east) in July 2024. On the same day, 4×100 mL aliquots were filtered by syringe through a custom-cut 25 mm ø, 0.45 µm PVDF filters (BioRad), housed in a syringe filter holder. The PVDF filter was folded and added to a 2 mL microcentrifuge tube and 380 µL ATL buffer and 20 µL Proteinase-K from a DNeasy Blood & Tissue Kit (Qiagen) was added. The filter was incubated at 56 °C overnight at 300 RPM on a Thermomixer C(Eppendorf) and then spun at 6000 RCF for 1 minute. The supernatant was collected and 200 µL of ethanol added. Following this, the sample was added to a DNeasy Mini spin column, and the DNeasy Blood & Tissue Quick-Start Protocol was followed from step 4. All collection glassware and syringe filter holders were bleached before use. See Fig S6 photos of the collection site and the sampled lake water.

### Sample Preparation and Amplicon Pooling

Environmental DNA (eDNA) was amplified with established primers targeting Fish 16S [22] species and Eukaryote 18S [23] rRNA genes. 16S priming pairs were, GACCCTATGGAGCTTTAGAC, CGCTGTTATCCCTADRGTAACT, and 18S priming pairs were gacatggttctaca-CACCGCCCGTCGCTACTACCG, cagagacttggtct-GGTTCACCTACGGAAACCTTGTTACG. The lowercase sections in the 18S primers are a fusion priming sequence not used in this study. PCR reactions were performed using GoTaq Hot Start Polymerase (Promega) at 0.1 U/µL, 5X Colorless GoTaq Flexi Buffer (Promega) at, Magnesium (Promega) at 4 nM, dNTPs (Promega) at 200 nM, Primers (IDT) at 500 nM. 2 µL DNA or 2 µL LTE (−ves) was added to the PCR master-mix to a total volume of 25 µL per reaction. PCR was performed with an initial denature step at 95° Cfor 5 min, followed by thermocycling with denature, annealing and extension steps of 95°C for 30 sec, 56°C for 30 sec, 72°C for 30 sec, respectively. A final extension step of 72°C for 10 min was also performed. Thermocycling was performed 40x for the 16S primers and 30x for the 18S primers.

Amplicon purification was performed through size selection bead-capture following manufacturer instructions for 2.5% w/v ‘Speed Beads’ magnetic carboxylate-modified particles (Sigma Alderich) and a custom capture buffer composed of 50% PEG (Sigma Alderich), 2.5 M NaCl (Sigma Alderich), 0.05% Tween-20 (Sigma Alderich), and 10 mM citric acid (pH 6) (Sigma Alderich).

For each extraction, equi-mass 16S and 18S amplicons (25 ng each) were pooled, resulting in approximately 200-400 pmol of total DNA, assuming amplicon lengths of 200-400 bp. Pooled amplicons were barcoded and prepared using the SQK-NBD114-24 kit (Oxford Nanopore Technologies), following the “VNBA_9168_v114_recO_15Sep2022” revision of the Native Barcoding Kit 24 protocol, which included a forward and reverse barcode specific to each extraction.

### Nanopore Sequencing

Sequencing was performed on a portable MinION Nanopore sequencing platform using the SQK-NBD114-24 native barcoding kit (Oxford Nanopore Technologies). The sequencing run lasted 21 hours, generating approximately 650,000 usable reads. Nanopore measurement parameters and read statistics are provided in Fig S8. Basecalling was conducted with Dorado 0.7.3 using the high accuracy model, without demultiplexing or trimming, resulting in a single FASTQ file. The command used was:

*dorado basecaller hac /pod5_pass_folder/ --kit-name SQK-NBD114-24 > 20240907_Pond_Scum_HAC-BC_No-Trimming*.*fastq --emit-fastq --no-trim*

Subsequent demultiplexing of the barcoded sequences was also performed using Dorado, which separated the sequences into individual FASTQ files and trimmed out the adapters and barcodes. Due to the short read lengths, no requirement was set for barcodes to be present at both ends of the sequence for demultiplexing. The command used was:

*dorado demux --kit-name SQK-NBD114-24 --output-dir /output-dir_folder/ --emit-fastq /fastq_folder/*

### FASTQ Reads Preprocessing and Analysis

FASTQ files from all barcoded samples were initially merged to create a comprehensive dataset. To ensure quality and relevance of the data, reads shorter than 100 base pairs (bp) and longer than 500 bp were removed from the dataset, as these lengths fall outside the expected range for the target amplicons and could represent sequencing artifacts or non-specific amplification.

To identify and quantify overrepresented sequences, the merged FASTQ files were processed using the ‘fastx_collapser’ tool, which clusters identical sequences. These read clusters were then further analyzed using custom Python scripts designed to remove sequences with high similarity, defined as those having three or fewer mismatches, or indels of two base pairs or less compared to another sequence within the cluster. This homology assessment was conducted by aligning the read clusters against themselves using BLAT (BLAST-like alignment tool), which allowed for precise identification of highly similar sequences.

Following the removal of highly similar reads, the remaining unique read cluster sequences were used to create an index with Bowtie2, a widely used tool for aligning sequencing reads to long reference sequences. The original FASTQ files were then mapped against the read cluster sequences using this Bowtie2 index, enabling the identification of reads that matched each cluster.

Coverage analysis was performed using Bedtools’‘genomecov’function, which calculates the depth of coverage for each sequence in the reference. This analysis provided insights into the proportion of 16S and 18S sequences that mapped to each read cluster, which was used as a proxy for estimating species abundance within the samples.

Finally, the read clusters were manually analyzed using NCBI nucleotide BLAST to identify the species of origin for each cluster. This step enabled the assignment of taxonomic labels to the read clusters, providing a detailed understanding of the species composition present in the environmental samples.

## Conclusions

The successful verification of the DIYNAFLUOR in both laboratory and field settings highlights its potential for supporting eDNA analysis in RLSs, in particular for Nanopore sequencing applications. Nonetheless, we acknowledge the difficulty some researchers may face in obtaining the required electronic components or 3D printed parts. While a full BoM and guidance for 3D printing is provided, we would recommend that potential users who are not confident in sourcing parts or 3D printing seek assistance from colleagues with relevant electrical or mechatronics engineering backgrounds. Future studies should also focus on improving the optical design to further increase the system’s overall sensitivity and accuracy, and we invite the open-source community to assist with this goal. However, at ∼2.6% the cost of the cheapest commercial alternative, we consider the stated accuracy, precision, sensitivity, and reproducibility a major achievement of the current DIYNAFLUOR platform. It is hoped that the DIYNAFLUOR facilitate further discussions regarding the democratization of molecular tools for eDNA analysis and demonstrates that open-source scientific equipment can provide access to current best-practice scientific methodologies.

## Acknowledgements

Author W. Anderson would like to thank the Australian Institute for Bioengineering and Nanotechnology, where he held an Adjunct Fellowship for the majority of this project, and the Trau Group, for providing laboratory access, reagents, and intellectual support that made the completion of this project possible.

