## Supplementary Information for "DIYNAFLUOR: An Affordable DIY Plug-and-Play Nucleic Acid Fluorometer for eDNA Quantification in Resource Limited Settings"

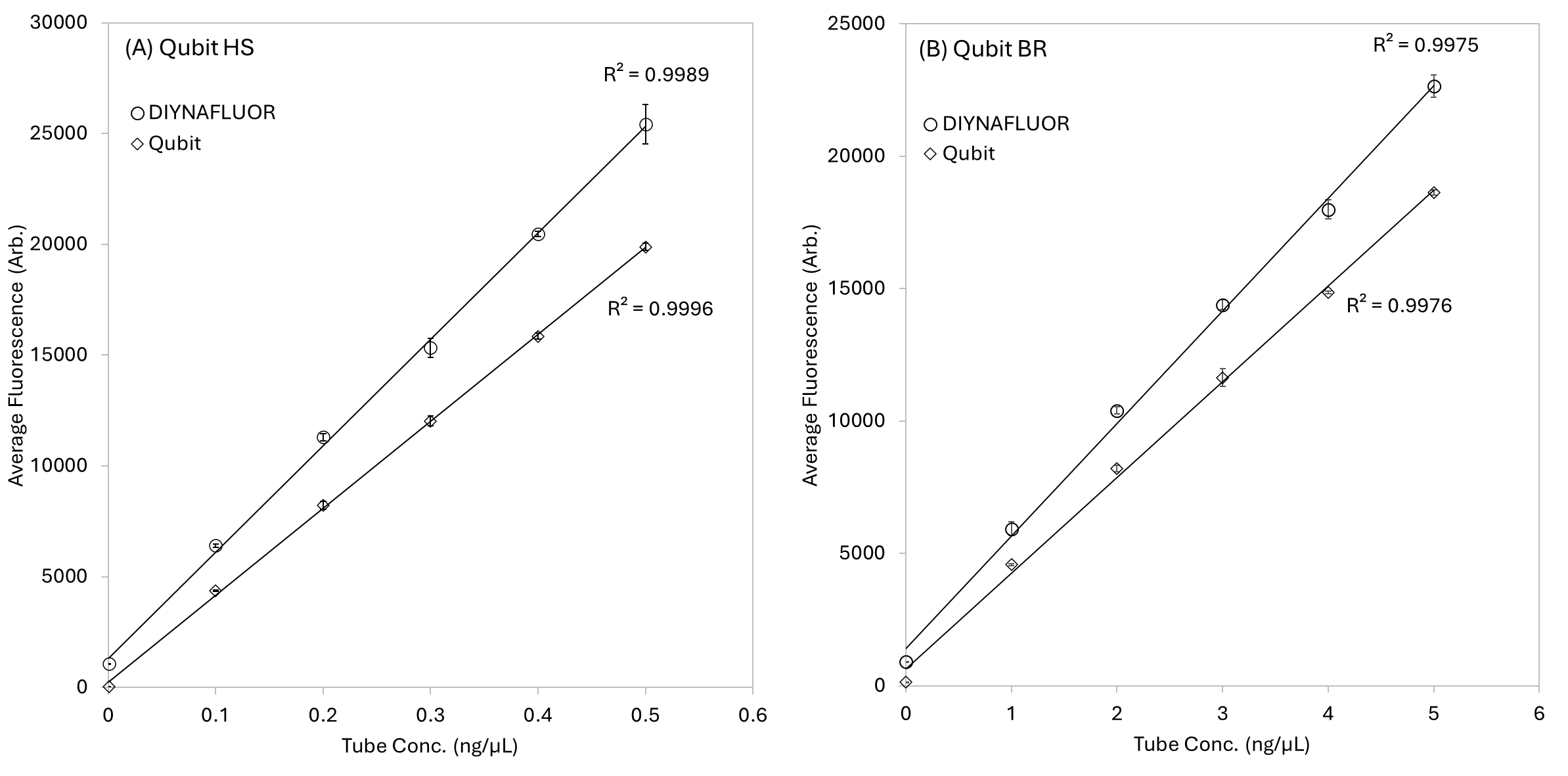

Figure S1. (A) Raw fluorescent data of 3n replicates of a quarter-fold dilutions series of DNA using the Qubit HS assay reagents. (B) Raw fluorescent data of 3n replicates of a quarter-fold dilutions series of DNA using the Qubit BR assay reagents.

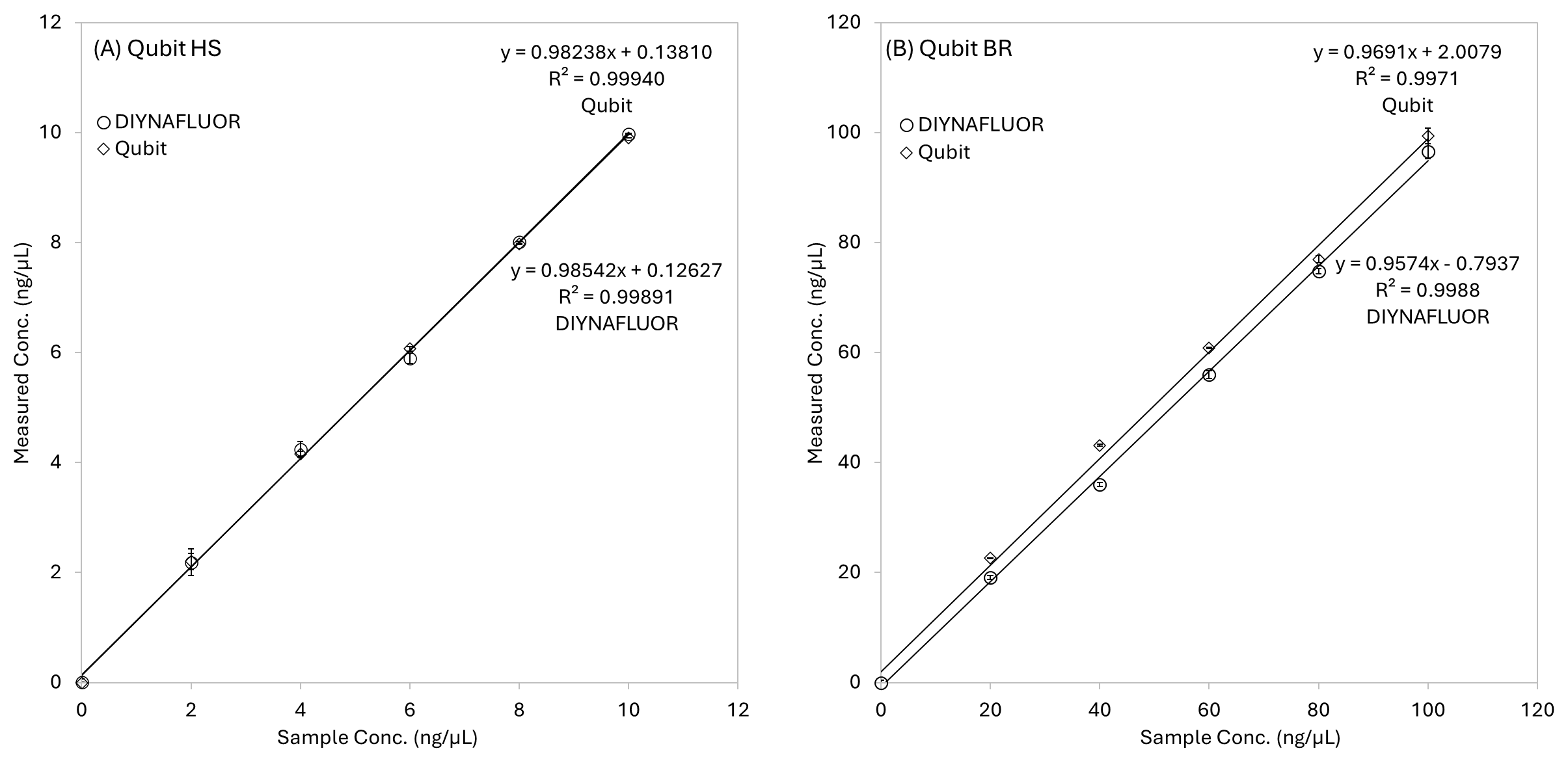

Figure S2. (A) Concentration measurements for 3n replicates of a quarter-fold dilution series of DNA using the Qubit HS assay reagents by both DIYNAFLUOR and Qubit 4. (B) Concentration measurements for 3n replicates of a quarter-fold dilution series of DNA using the Qubit BR assay reagents by both DIYNAFLUOR and Qubit 4.

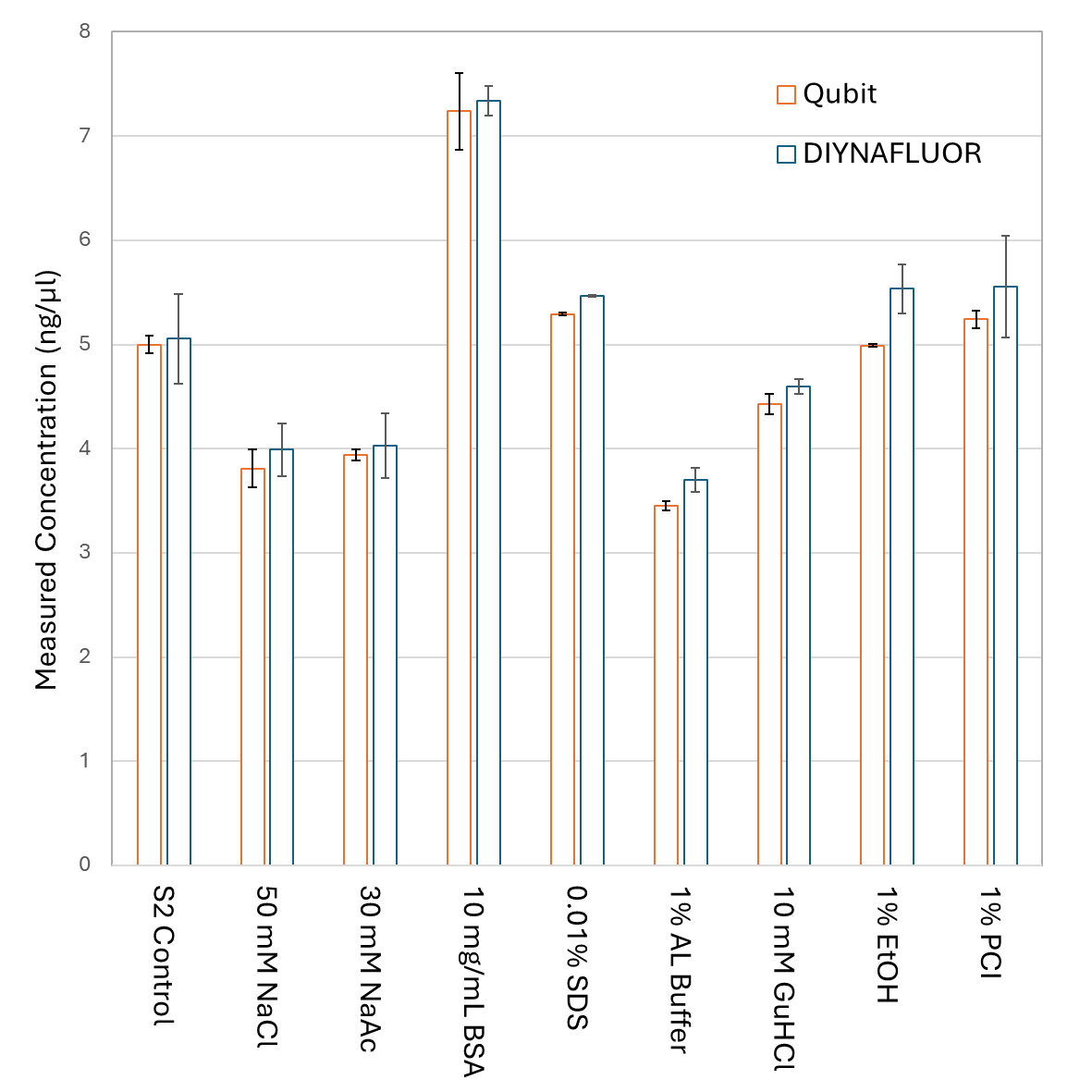

Figure S3. Robustness assay for contaminants that may carry-over in common DNA extraction protocols. 2n replicate measurements were performed by both DIYNAFLUOR and Qubit 4 using the Qubit HS DNA assay kit with contaminants spiked into DNA at 5 ng/µL. In general, deviation from the expected value of 5 ng/µL correspond similarly between the DIYNAFLUOR and Qubit measurements. However, these result, which deviate from the expected value by greater than 40% in the case of the BSA test, highlight the importance of optimising extraction protocols for DNA purity for accurate fluorometric results.

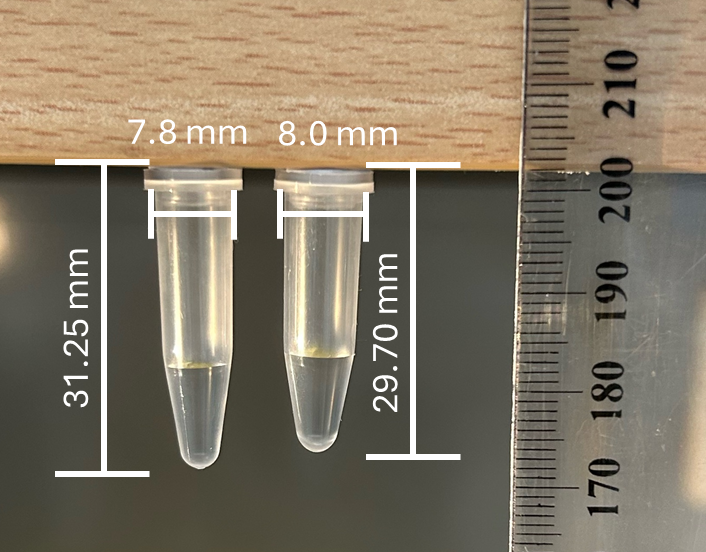

Figure S4. The two different conformations of 0l.5 mL PCR tubes we have identified being supplied with Qubit kits. Using different tube geometries interchangeable in the same measurement can result in unexpected deviations due to the different optical pathlengths. Wider tubes can also form a snug fit in the Sample Well if printing tolerances are off, which can lead to users not fully inserting tubes for measurement. The Axygen’s 0.5 mL thin-walled PCR tubes used in the SYBR-Safe assay (not pictured), have a thickness similar to the tube on the left.

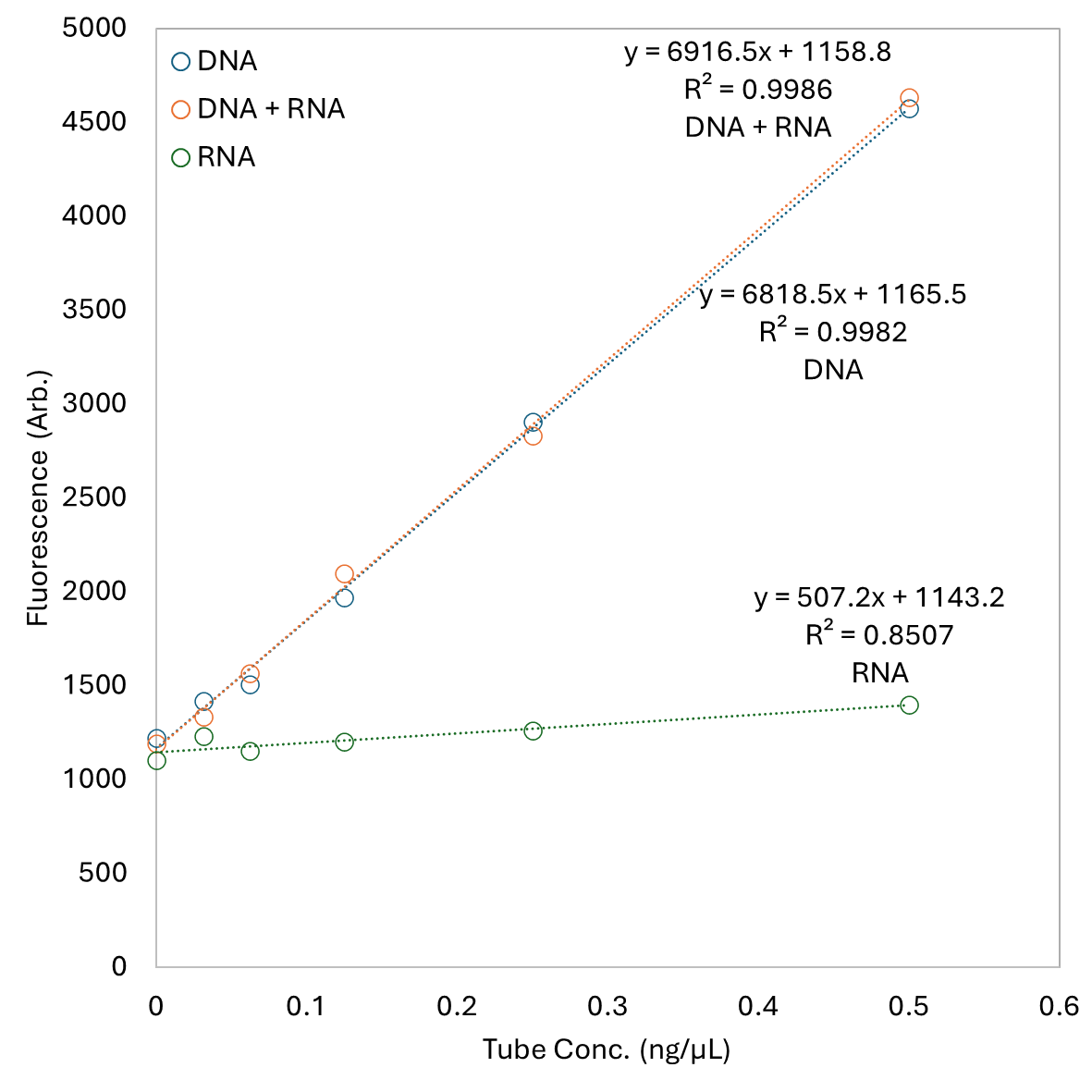

Figure S5. Due to many DNA purification protocols also recovering RNA, a robustness assessment of the SYBR-Safe assay for RNA sensitivity was performed. Measurements were performed on DNA, DNA+RNA and RNA dilution series. SYBR-Safe was shown to have a significantly lower fluorescent response to RNA, and was not seen to significantly affect the DNA+RNA results compared to the DNA-only measurement.

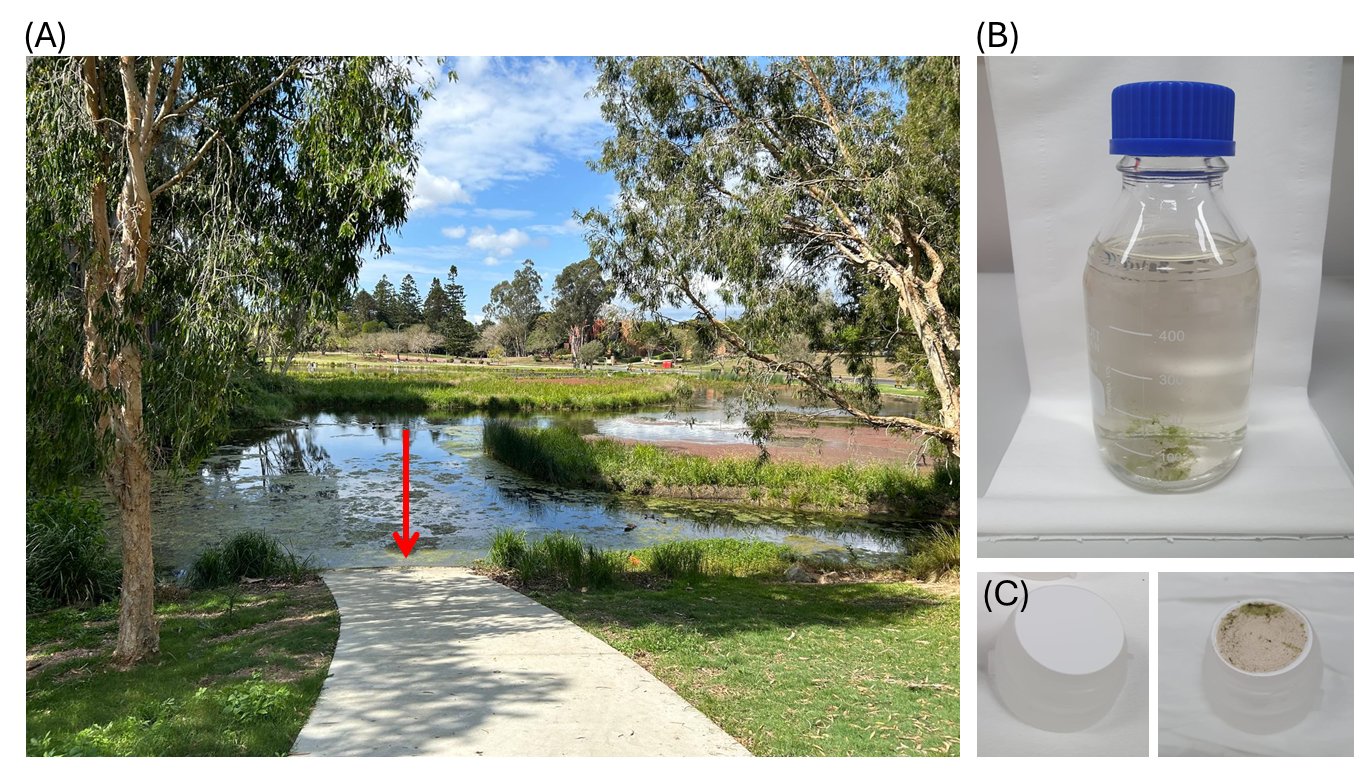

Figure S6. (A) The location at The University Queensland, St Lucia, Australia where lake water was collected. (B) The collected lake water. The Schott bottle was bleach decontaminated bedfore collection. (C) Before and after example of a 0.45 mm PVDF filter membane in a syrynge filter holder that had 100 mL of lake water filtered through it. The syringe filter holder was bleach decontaminated before use.

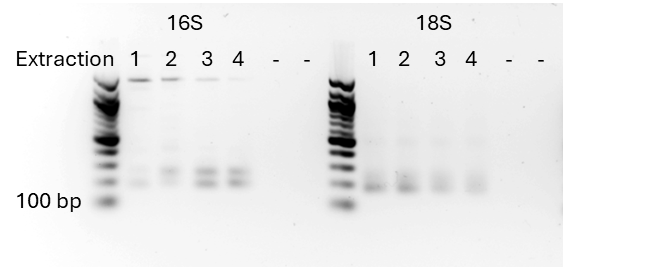

Figure S7. Gel electropherogram of the 16S and 18S amplicons against a 100 bp ladder. Expected product size was 200-300 bp. Some large off target amplicon were observed in the 16S samples but were excluded from sequencing analysis.

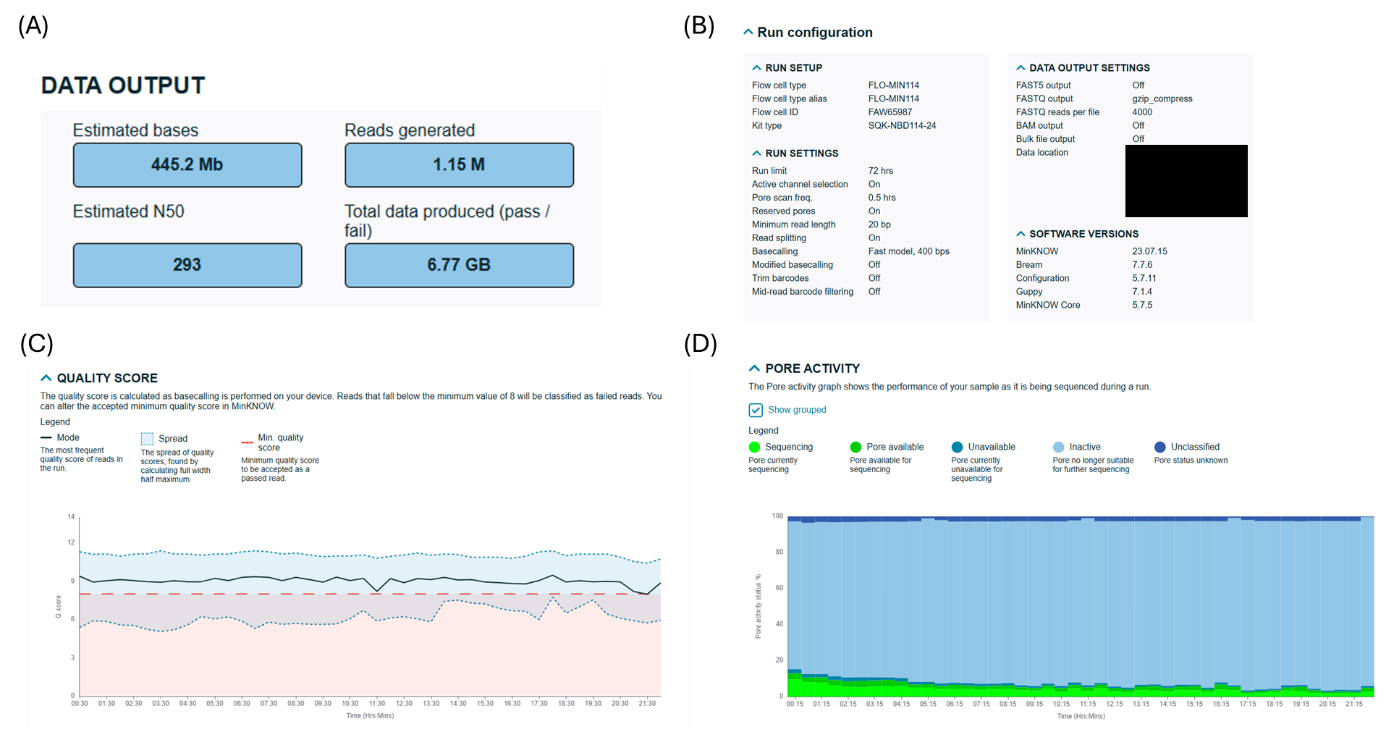

Figure S8. (A) Statistics for the nanopore sequencing run of the 16S and 18S metabarcode libraries. (B) Nanopore sequencing run parameters. (C) Quality metrics over the 21.5-hour sequencing run. (D) Pore activity over the 21.5-hour sequencing run. A pre-used nanopore flow cell, with only ~100 remaining active pores, was used for this study to demonstrate that heavily degraded flow cells can still be effective in RLSs. The Flow cell was DNAse treated before reuse (following Oxford Nanopores protocol) to remove DNA contamination from previous sequencing runs.

Table S1. The 45 identified species from Nucleotide Blast of the identified from the sequence clustering bioinformatics pipeline analysis of the nanopore reads.

| **Sequencing ID** | **Sequence Length** | **Extraction 1** | **Extraction 2** | **Extraction 3** | **Extraction 4** | **GenBank ID** | **Scientific Name** |
| --- | --- | --- | --- | --- | --- | --- | --- |
| 1-548 | 181 | 7582 | 5452 | 14560 | 26860 | DQ470581.1 | Lathonura rectirostris |
| 2-503 | 171 | 7013 | 8877 | 3582 | 5707 | OU230632.1 | Halteria sp. |
| 6-264 | 245 | 3 | 10952 | 13503 | 20177 | KF665157.1 | Rhinella marina |
| 13-161 | 104 | 147 | 510 | 91 | 39 | PP923469.1 | Trichosurus vulpecula |
| 17-140 | 178 | 6349 | 4168 | 3053 | 3213 | MW775240.1 | Eocercomonas sp |
| 37-56 | 178 | 801 | 2829 | 4385 | 3059 | MW075316.1 | Pediastrum duplex |
| 50-34 | 175 | 1118 | 332 | 1465 | 1383 | LC647555.1 | Cryptomonas borealis |
| 61-29 | 180 | 1316 | 1435 | 768 | 965 | AB749074.1 | Uncultured chrysophyte |
| 62-29 | 178 | 3571 | 2624 | 6935 | 14089 | DQ470581.1 | Lathonura rectirostris |
| 64-28 | 180 | 2370 | 2195 | 1704 | 2141 | EF060630.1 | Dothioraceae |
| 67-27 | 207 | 165 | 153 | 908 | 9521 | AM490291.1 | Pleuroxus truncatus |
| 70-26 | 173 | 1219 | 1279 | 842 | 1066 | EU143957.1 | Uncultured cryptophyte |
| 71-25 | 189 | 338 | 190 | 60 | 37 | CP011834.1 | Limnohabitans sp. |
| 75-25 | 178 | 140 | 401 | 1686 | 2124 | MN696693.1 | Nitzschia paleaeformis |
| 78-25 | 178 | 2355 | 2603 | 2278 | 2747 | AY651083.1 | Pedospumella encystans |
| 79-24 | 186 | 1 | 832 | 3341 | 2797 | KY273259.1 | Netzelia tuberculata |
| 82-22 | 100 | 113 | 32 | 174 | 51 | AJ244830.1 | Anguilla reinhardtii |
| 86-21 | 180 | 1642 | 1325 | 675 | 840 | NG_063470.1 | Filobasidium uniguttulatum |
| 91-20 | 182 | 601 | 694 | 714 | 866 | XR_005548808.1 | PREDICTED: Eucalyptus grandis |
| 97-19 | 182 | 6 | 10 | 1 | 1544 | AY821866.1 | Dicrotendipes fumidus |
| 109-18 | 180 | 767 | 791 | 543 | 605 | EF060945.1 | Tremellales sp |
| 113-17 | 172 | 161 | 158 | 656 | 855 | L28811.1 | Chilomonas paramecium |
| 120-16 | 170 | 3901 | 4465 | 1904 | 2616 | HQ219432.1 | Uncultured ciliate |
| 122-16 | 183 | 77 | 79 | 43 | 1606 | AJ012519.1 | Stenostomum leucops |
| 125-16 | 172 | 134 | 215 | 383 | 1388 | KY019066.1 | Sarcocystis silva |
| 127-16 | 157 | 270 | 58 | 432 | 329 | Many Species |  |
| 128-15 | 180 | 2465 | 2746 | 1621 | 2084 | KY947997.1 | Umbilicaria muehlenbergii |
| 139-14 | 176 | 508 | 557 | 488 | 533 | KX442736.1 | Poteriospumella sp. |
| 145-14 | 184 | 285 | 329 | 629 | 195 | EU853660.1 | Arachnula impatiens |
| 158-13 | 180 | 644 | 567 | 430 | 549 | KY981682.1 | Myrmecia israelensis |
| 164-13 | 180 | 2328 | 2140 | 1355 | 1465 | AF314999.1 | Bullera pseudoschimicola |
| 166-13 | 170 | 1083 | 1424 | 568 | 957 | HQ219432.1 | Uncultured ciliate clone |
| 168-13 | 173 | 187 | 134 | 228 | 268 | LC647555.1 | Cryptomonas borealis |
| 172-13 | 169 | 2344 | 2898 | 1143 | 1791 | EF024884.1 | Uncultured Dunaliellaceae |
| 177-13 | 178 | 166 | 530 | 1191 | 2255 | FN398345.1 | Navicula sp. |
| 185-12 | 180 | 953 | 697 | 1871 | 3084 | DQ470581.1 | Lathonura rectirostri |
| 188-12 | 175 | 355 | 321 | 394 | 387 | LC647557.1 | Cryptomonas sp. |
| 190-12 | 176 | 422 | 518 | 375 | 433 | KX431464.1 | Chrysophyceae sp. |
| 196-11 | 154 | 104 | 255 | 373 | 561 | KF524428.1 | Vorticella sp. |
| 198-11 | 180 | 217 | 223 | 372 | 375 | HQ219364.1 | Uncultured stramenopile clone |
| 201-11 | 171 | 296 | 676 | 139 | 204 | FJ543106.1 | Strombidium sp. |
| 207-11 | 178 | 104 | 274 | 928 | 1415 | AJ867277.1 | Nitzschia amphibia |
| 213-11 | 182 | 221 | 865 | 1448 | 1507 | MT840106.1 | Keratella serrulata |
| 220-10 | 180 | 1427 | 1256 | 819 | 948 | FO905468.1 | Leptosphaeria biglobosa brassicae |
| 221-10 | 180 | 883 | 656 | 434 | 500 | KY101795.1 | Papiliotrema pseudoalba |
| 232-10 | 231 | 0 | 183 | 201 | 0 | No Result |  |

Table S2. Nanopore read statistics for each of the extraction 16S + 18S amplicon libraries. Total Reads were 205 k ± 46 k across the four libraries, indicating the DIYNAFLUOR-based molarity calculations were able to provide equivalent read coverage from the 4 samples. Deviations are most likely due to the different amplicon product sizes generated, as shown in Figure S7.

| **Barcode** | **Total bases (Mb)** | **Passed bases (%)** | **Total reads (k)** | **Passed reads (%)** |
| --- | --- | --- | --- | --- |
| Extraction 1 | 60.375475 | 77.6 | 161.672 | 79.7 |
| Extraction 2 | 65.35377 | 77.8 | 190.391 | 79.4 |
| Extraction 3 | 62.081334 | 79.1 | 201.886 | 80.8 |
| Extraction 4 | 79.360908 | 77.3 | 269.574 | 79.2 |

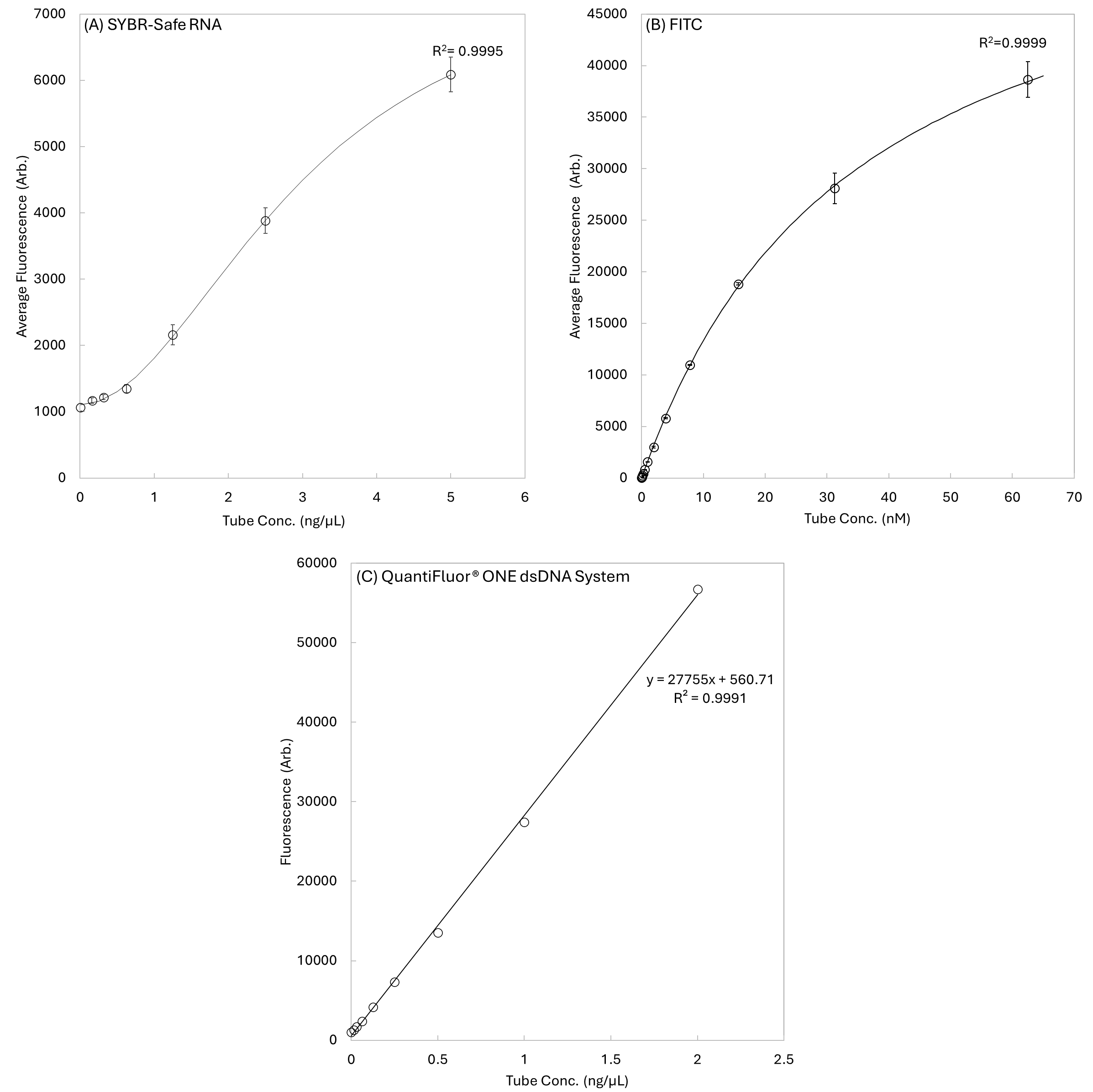

Figure S9. (A) Fluorescent response of a 2-fold dilution series of RNA using the low-cost SYBR-Safe assay (3n replicates). The working range of this assay is an order of magnitude larger than the input RNA concentrations used in the DNA robustness assay in Figure S5. (B) Fluorescent response of a 2-fold dilution series of FITC (3n replicates). A modified non-linear least-squares it of a Hill Equation was used to fi the data in (A) and (B). (C) A 2-fold dilution series of the commercial Quantifluor ONE dsDNA system (Promega), showing a similar fluorescent response to DNA input concentration as the Qubit HS DNA assay.

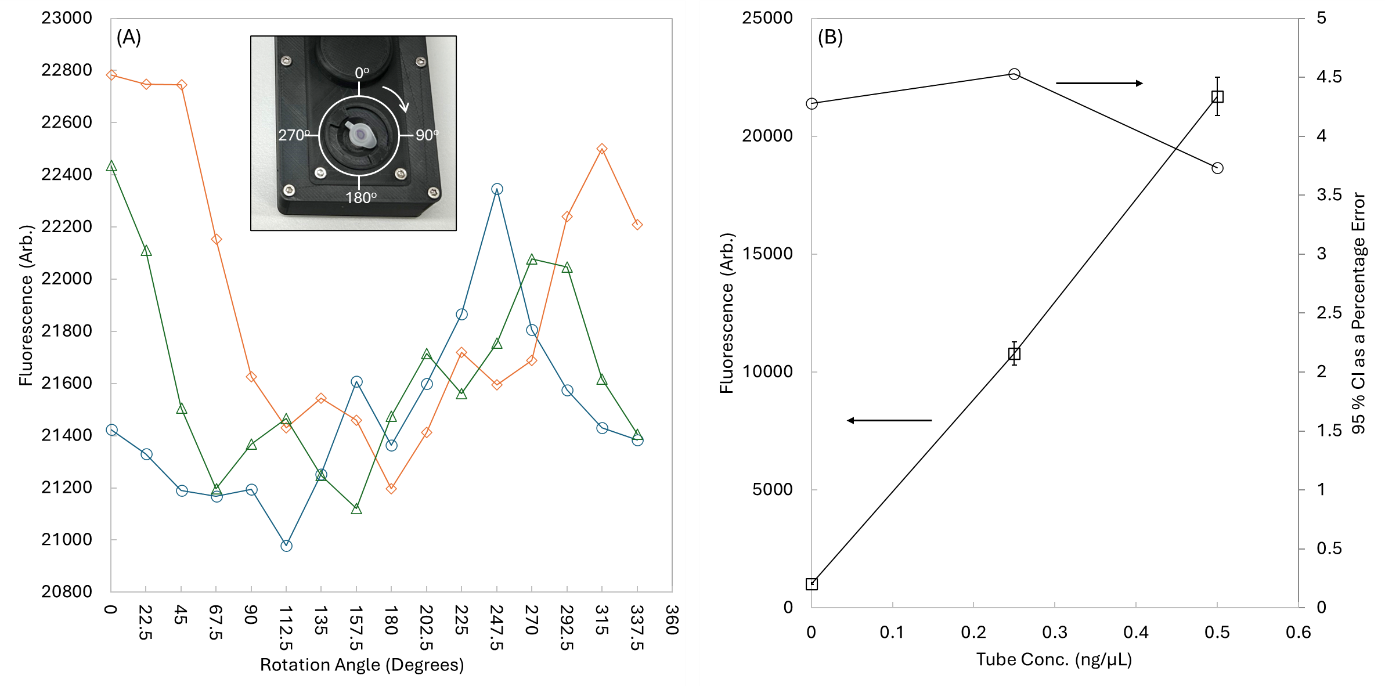

Figure S10. (A) Fluorescent output of ‘in-assay’ DNA concentration of 0.5 ng/µL using the Qubit HS reagents (3n replicates) measured at 22.5-degree rotation points. The inset figure demonstrates how the measurements were made. These data show that changes in the optical pathlength from factors like inconsistencies in the tubes optical profile around its vertical axis, and small deviations in placement of the tube in the sample well, are the major cause of random measurement error in the DIYNAFLOUR system. (B) A statistical analysis of random noise in the DIYNAFLOUR system cause by tube placement. The left axis shows the fluorescent signal for 3n replicates of measurements performed with DNA concentrations 0, 0.25 and 0.5 ng/µL using the Qubit HS DNA reagents, across 22.5 degree rotation measurements (as demonstrated in (A)), error bars represent the 95% CI in range of fluorescent response. The right axis shows the 95% CI as a percentage error for the fluorescent signal, showing that random error in measurements is 4.5% or lower across the working range of the DIYNAFLUOR system for this assay. It should be noted that bleaching of the fluorophores over repeat measurements will result in these error values being larger than expected, however we believe this value to be the truest current measure of random error in the DIYNAFLUOR system. It should be noted that this error is smaller than the acceptable error of 8% when pipetting 1 µL with a 10 µL pipette, as defined by ISO 8655.
